# Multivariate Analysis of PET Pharmacokinetic Parameters

**DOI:** 10.1101/2022.05.04.490593

**Authors:** Granville J. Matheson, R. Todd Ogden

## Abstract

**Purpose:** In positron emission tomography (PET) quantification, multiple pharmacokinetic parameters are typically estimated from each time activity curve. Conventionally, all but the parameter of interest are discarded before performing subsequent statistical analysis. However, we assert that these discarded parameters also contain relevant information which can be exploited to improve the precision and power of statistical analyses on the parameter of interest. Properly taking this into account can thereby draw more informative conclusions without collecting more data.

**Methods:** By applying a hierarchical multifactor multivariate Bayesian approach, all estimated parameters from all regions can be analysed at once. We refer to this method as PuMBA (Parameters undergoing Multivariate Bayesian Analysis). We simulated patient-control studies with different radioligands, varying sample sizes and measurement error to explore its performance, comparing the precision, statistical power, false positive rate and bias of estimated group differences relative to univariate analysis methods.

**Results:** We show that PuMBA improves the statistical power for all examined applications relative to univariate methods without increasing the false positive rate. PuMBA improves the precision of effect size estimation, and reduces the variation of these estimates between simulated samples. Furthermore, we show that PuMBA yields performance improvements even in the presence of substantial measurement error. Remarkably, owing to its ability to leverage information shared between pharmacokinetic parameters, PuMBA even shows greater power than conventional univariate analysis of the true binding values from which the parameters were simulated. Across all applications, PuMBA exhibited a small degree of bias in the estimated outcomes, however this was small relative to the variation in estimated outcomes between simulated datasets.

**Conclusion:** PuMBA improves the precision and power of statistical analysis of PET data without requiring the collection of additional measurements. This makes it possible to study new research questions in both new and previously collected data. PuMBA therefore holds great promise for the field of PET imaging.

## 1 Introduction

Positron emission tomography (PET) is an *in vivo* neuroimaging method with high biochemical sensitivity and specificity. It is an essential tool for the study of the neurochemical pathophysiology of psychiatric and neurological disease, as well for pharmaceutical research. However, PET is a very costly and invasive procedure that involves exposing participants to radioactivity, thereby limiting the feasibility of large studies. As a result, low statistical power is a common obstacle encountered for studying clinically relevant research questions. Efforts to improve the power of PET imaging have typically focused on the development of new radiotracers with improved sensitivity as well as new pharmacokinetic (PK) models with greater accuracy; more recently there have been data standardisation and sharing initiatives to foster inter-group collaboration and increase sample sizes [1–3]. However, there has been comparatively little attention paid to the development of more nuanced statistical analysis of PET data for the same purpose.

PET quantification involves fitting PK models to a series of radioactivity concentrations in a region of the brain over time, called a time activity curve (TAC), most often using nonlinear least squares (NLS) optimisation. These models typically consist of between 1 and 5 parameters of which one (or a function of two or more parameters) is used as measure of the binding of the radioligand to the target protein. Once the TAC data from all regions and all subjects have been fit using the selected model, the parameter estimates reflecting target binding for each region and subject are then entered into a subsequent statistical model, e.g., a *t*-test comparing patients and control subjects, while the other estimated parameters are not taken into account in the analysis.

We recently introduced SiMBA [4], which makes use of Bayesian hierarchical multifactor modelling to fit PET TAC data and perform statistical analysis simultaneously across both individuals and regions. The primary disadvantage of this technique is that it is highly computationally intensive, and currently only implements the two-tissue compartment model [5]. However, the model improves the estimation of binding parameters, and yields substantial advantages in terms of increased precision and statistical power for statistical comparisons. Intriguingly, in simulation studies we found that the statistical power of SiMBA for detecting group differences was even greater than for univariate statistical analysis performed on the “true” binding values from which the TACs were generated. This suggests that even if binding measures could be measured exactly for each subject, it would still not be possible to attain the statistical power that we observed with SiMBA.

Upon further inspection, we discovered that this performance gain could be explained by the multivariate modelling strategy employed in SiMBA. In other words, instead of extracting only a single parameter as a measure of binding, statistical analysis was performed using all estimated parameters simultaneously, thereby allowing the model to exploit shared information among all the PK parameters. This general concept is demonstrated in Figure 1: the shape of the 2-dimensional density plot of the two variables is highly dependent upon their correlation with one another. If two parameters are highly correlated with one another, then the conditional variance of each parameter at any given value of the other is considerably smaller. Hence, if the estimation of both variables and their correlation with one another are sufficiently precise, then the *conditional* variance of estimated parameters can be reduced to below that of the marginal true values. In other words, by exploiting shared information between parameters, even if those parameters are not directly relevant to the statistical contrast of interest, the performance of the statistical model can be improved.

**Fig. 1.**
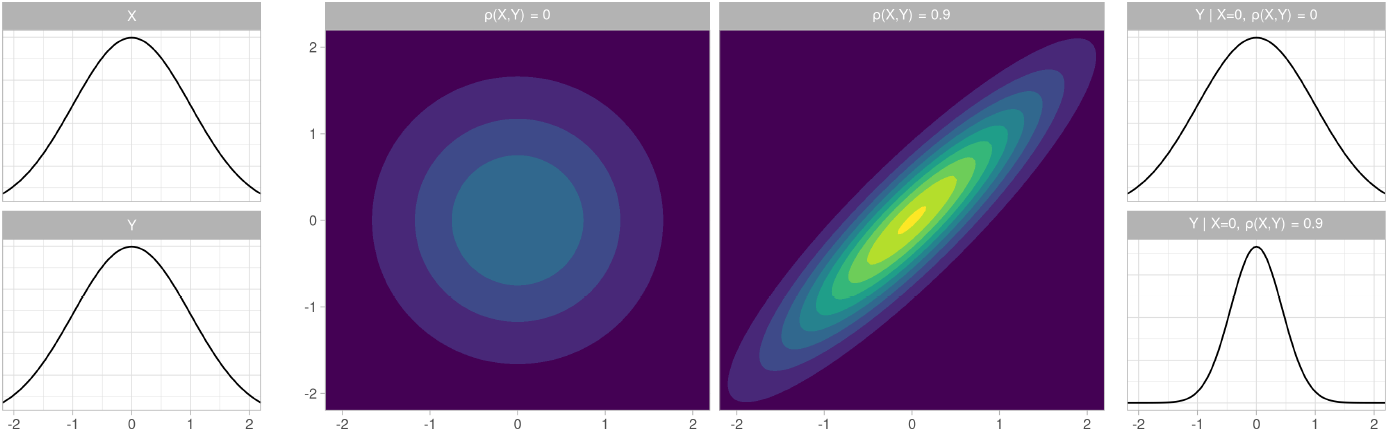
Comparisons of marginal and conditional densities for multivariate normal distributions. Left: Marginal densities of variables X and Y after standardisation. Middle: Multivariate contour plots of the two-dimensional densities of X and Y, with either no correlation or a strong correlation between them. Right: Conditional densities of variable Y conditional on X when there is either no correlation or a strong correlation between variables X and Y.

In this study, we evaluate whether applying a multivariate statistical analysis to PET pharmacokinetic outcome parameters estimated in the conventional manner using NLS estimation can also provide inferential advantages, without needing to fit the full SiMBA model to the dynamic TAC data. We refer to this approach, by analogy with SiMBA, as Parameters undergoing Multivariate Bayesian Analysis: PuMBA. The computational requirements for this modelling strategy is on the order of minutes on a single core, compared with days for SiMBA, and can readily be adapted to a wider range of pharmacokinetic models, thereby facilitating its application to a broader range of research questions. PuMBA may therefore serve as a convenient intermediate substitute for a full SiMBA analysis.

## 2 Methods

### 2.1 Model Specification

PuMBA can be described as a multivariate hierarchical multifactor model. It is multivariate in that there are multiple dependant variables estimated at once — in contrast with a multivariable model in which there are multiple *independent* variables. It is hierarchical in that it makes use of “partial pooling”. This means that parameters are modelled as originating from a common distribution, and are therefore shrunk towards the global mean in an adaptive regularisation process. This shrinkage allows the model to take advantage of similarities between individuals within the dataset to improve its inferences [6–8]. Finally, PuMBA is multifactor in that there are multiple hierarchies at once within which we perform partial pooling [9].

For PuMBA, as for SiMBA, linear models are defined for each of the PK parameters, defined by an intercept, covariates, and partially pooled deviations from the expectation value for each individual *j* and region *k*, for each of the *m* PK parameters. We define a global mean intercept (*α*) for each parameter, representing the mean value for that parameter. For each PK parameter *i*, the influence of covariates of individual *j* are expressed by a covariate vector (*β_i_*) multiplied by a covariate matrix (***X**_i,j_*). These covariate matrices are defined independently for each PK parameter, and can include variables such as age, sex, or group membership. Lastly, we define an additive sequence of differences for each of the separate hierarchies [9]: across individuals (*τ_j_*), across regions (*ν_k_*), as well as a final term for residual variation (*ϵ_j,k_*).

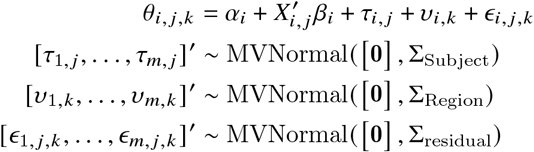

This defines the generalised model framework, in which estimation is performed using partial pooling for all parameters across all hierarchies, resulting in some degree of shrinkage towards the mean. In practice, shrinkage of most parameters towards a shared mean is desirable, however regional differences in certain parameters are so heterogeneous that a common distribution cannot be assumed owing to regional neuroanatomical differences. For this reason, blood delivery and binding parameters are estimated independently from one another without pooling, i.e., using fixed effects. More details are provided in the following section.

### 2.2 Model Implementation

Firstly, all PK parameters are transformed to their natural logarithms. This serves several purposes. Firstly, this naturally constraints all parameters to be positive, corresponding to their theoretical range as biological quantities and rate constants. Secondly, this serves to define additive differences within the linear model as proportional differences in the original quantity, since biological differences or changes in PET are typically assumed to exhibit similar proportional, as opposed to absolute, differences between different regions or individuals. Lastly, this serves to stabilise the variance between regions: in PET, we typically make the assumption that the proportional variance between regions is relatively similar.

The input parameters for PuMBA are the PK parameters estimated by the kinetic model from the TACs using NLS. For the two-tissue compartment (2TC) model [10], the model parameters were ***K***_1_, ***V***_ND_, ***BP***_P_, and *k*_4_. For the 1TC [10], we used **K**_1_ and ***V***_T_. Finally, for the SRTM [11], we used 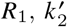 and ***BP***_ND_. We selected parameterisations of the model parameters in such a way as to improve the ease by which priors could be defined. To this end, for each model we defined a binding parameter and a blood delivery parameter, and defined the remaining PK parameters in such a way as to maximise the extent to which shrinkage towards a common mean value is most theoretically motivated, i.e. which can be considered as originating from a common distribution. For the 2TC, ***BP***_P_ was selected over ***BP***_ND_, as the former is more identifiable using NLS and in simulations, estimated values show stronger correlations with the true values compared to ***BP***_ND_. For SRTM, we used 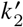 rather than ***k***_2_ as the former parameter is a property of the reference region and should theoretically be fairly consistent between regions within each individual, similar to how this parameter is set to a global estimate using SRTM2 [12].

For the definition of covariate matrices for each parameter, this task can depend both on the tracer as well as the sample itself. For instance, age might be a predictor for both blood delivery and binding. On the other hand, patient status might only be included as a predictor for binding – unless the condition is also thought to affect regional blood delivery, in which case patient status might also be included as a predictor for blood delivery. Careful judgement should be applied to this task, although model comparison methods can also be helpful [6, 13, 14].

### 2.3 Model Fitting

We make use of multivariate Bayesian hierarchical multifactor modelling to fit the model described above using Markov Chain Monte Carlo (MCMC) sampling. We defined the model using the STAN probabilistic programming language [15], which applies Hamiltonian Monte Carlo (HMC), with code generated using `brms` 2.15.0 [16] using R version 4.0.5 *(Shake and Throw*) [17].

Priors were specified in such a way as to exclude parameter values which could be deemed as unlikely *a priori* based on domain knowledge, but not to greatly inform the model. We used moderately informative normal priors for the intercept (**α**) terms, and zero-centred half-normal regularising priors for the standard deviation of all pooled parameters. LKJ [18] priors were defined for correlation matrices. More details are provided in Supplementary Materials S1.

NLS parameter estimation was performed using `kinfitr` [19, 20] for the one-tissue compartment model (1TC) and the simplified reference tissue model (SRTM) [11]. For the two-tissue compartment model (2TC), the model was fitted directly using NLS using an analytical convolution of the arterial input function with the impulse response function, as previously described [4], solving for ***K***_1_, ***V***_ND_, ***BP***_P_ and ***k***_4_. In all cases, weights were estimated using the default kinfitr weighting scheme.

### 2.4 Simulations

For the purpose of assessing the performance of this modelling approach, we generated simulated datasets to compare the proposed methodology with that of the conventional approach. Simulation parameters were generated based on the posterior mean values of parameters estimated from empirical data by fitting the relevant model to the data and simulating from the estimated parameters. The datasets used were as follows: 97 individuals measured with [^11^C]WAY100635 [21], 16 measurements from 8 individuals measured with [^11^C]ABP688 [22], 47 measurements with [^11^C]DASB from 33 individuals [23, 24], and 23 individuals measured with [^11^C]GR103545 [25]. Simulated datasets had between 10 and 100 individuals in each of a patient and a control group, i.e. between 20 and 200 individuals in total, with between 8 and 9 regions for each ligand (more details in Supplementary Materials S2).

We simulated new sets of individuals by simulating from the estimated multivariate and univariate normal distributions describing variation between individuals and regions within individuals. When simulating regional variation, we used the estimated values for each region, rather than simulating from the estimated distributions. In this way, we simulate from the same set of regions, but within a unique set of individuals, with a unique set of individual variations at the regional level.

We set global group differences in the natural logarithm of the binding parameter to be equal to 0.1, corresponding to a 10.5% difference between groups, and separately to zero to assess the false positive rate. Simulated data were generated without any effects of age or sex, and so these covariates were not included in the PuMBA models applied to the simulated data. The univariate analyses performed included *t*-tests and linear mixed effects (LME) models with the natural logarithm of the parameter representing ligand binding as the dependent variable, considering group and region as fixed effects and subject as the only random effect. The *t*-tests were performed independently for each region of the dataset. LME analysis was performed across all regions using `lme4` [26].

We evaluated power and false positive rates by fitting logspline density functions [27] to the upper and lower bounds of the 95% confidence/credible intervals of the estimated difference between groups. We then assessed the cumulative density of these fitted density functions above and below zero, which we could use to estimate the proportion of simulated datasets for which a the 95% confidence/credible interval would overlap with (or not overlap with) zero. We have shown previously that this method closely aligns with empirical estimates, and allows for the estimation of power in small numbers of trials [4].

#### 2.4.1 TAC Simulations

For TAC simulations, we first fit the TAC data using SiMBA, and then used the posterior mean values of the SiMBA fit to simulate new TAC data. To model these data, the TACs were first fitted using NLS to generate PK parameters, which served as the input to the LME and PuMBA models. Data were generated with the standard deviation of the measurement error equal to that estimated in the original sample. We also considered half, double and quadruple the original measurement error to assess the sensitivity of the different approaches to the magnitude of the measurement error. We assessed the performance of these approaches using 500 simulated datasets for each condition.

We also assessed the performance of PuMBA in the same simulated TAC datasets for each condition as examined previously with SiMBA [4], in order to compare the performance of PuMBA and SiMBA in the same data. In this dataset, group differences were equal to 0.182 in the natural logarithm of ***BP***_P_, corresponding to a group difference of 20%. This included 50 simulated datasets for each condition owing to the much-greater computational burden of SiMBA.

The simulation parameters are provided in Supplementary Materials S2. The binding parameter for which group differences were simulated and estimated, was ***BP***_P_.

#### 2.4.2 Parameter Simulations

While the TAC simulations above were based on parameters estimated using SiMBA, SiMBA only currently applies the 2TC. For SRTM and 1TC, we therefore simulated data from estimated PuMBA parameters instead. To this end, the empirical TAC data were first fitted using NLS to generate PK parameters, which were then modelled using PuMBA. The posterior mean values of the PuMBA model fit were used to simulate new sets of parameter estimates. PuMBA parameters, however, do not allow for the discrimination of variance originating from error in the estimation of the PK parameters using NLS, and variation at the region-within-individual level: both of these sources of variation are part of the residual variance, ***ϵ***. For this reason, simulating TAC data from PuMBA parameters and subsequently estimating PK parameters from these TACs using NLS would effectively amount to doubling the influence of measurement error. It would be present firstly in the estimated ***ϵ*** matrix from the PuMBA model fitted to the empirical data, as well as from the estimation of the PK parameters from the generated TACs themselves. For this reason, for SRTM and 1TC, we estimated only parameter data from the PuMBA parameters, and not full TACs from these parameters. To model these data, PuMBA, LME and *t*-tests were applied to the simulated parameter estimates, using 250 simulated datasets for each condition.

We made use of data from two radioligands for each model: for the one tissue compartment (1TC) model, we used [^11^C]DASB and [^11^C]GR103545; and for the simplified reference tissue model (SRTM), we used [^n^C]WAY100635 and [^11^C]DASB with cerebellar white and grey matter respectively as reference region corresponding with previous recommendations [28–31]. The simulation parameters are described in Supplementary Materials S2. The binding parameter used for each model in which group differences were simulated and estimated, were ***BP***_ND_ for SRTM and for the 1TC model.

## 3 Results

### 3.1 TAC Simulations

For the TAC simulations, we compare the performance of PuMBA to that of LME models applied to the estimated ***BP***_P_ values. For comparison, we also applied LME to the “true” ***BP***_P_ values from which the TACs were simulated, i.e. representing the “ideal” case in which binding parameters are perfectly estimated, incorporating only individual-level and regions-within-individual-level variation. The results of these simulations are shown in Figure 2. Naturally, we see that the power of both LME and PuMBA are decreased with increasing measurement error, although these decreases are of less consequence for LME compared to PuMBA. Although concerns have previously been raised about the accuracy of direct quantification of ***BP***_P_ [32], i.e. without the use of a reference region, we observe that LME applied to the estimated ***BP***_P_ values with the original measurement error exhibits similar or only marginally reduced power compared to when LME is applied to the true values. This supports the use of ***BP***_P_ as a sufficiently good index of specific binding for these two tracers.

**Fig. 2.**
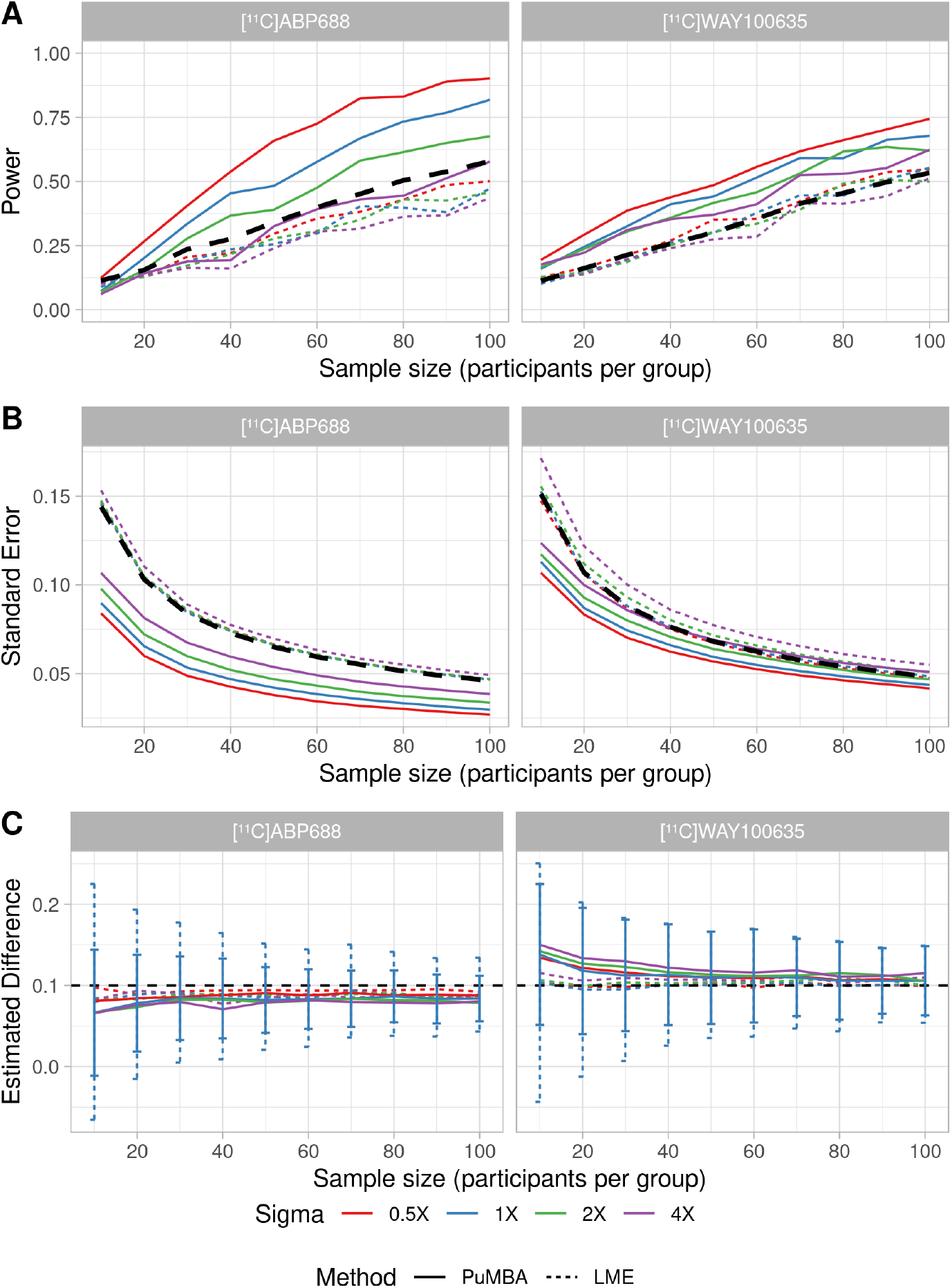
TACs were simulated with the standard deviation of measurement error equal to the same, half, double or quadruple the estimated measurement error in the original data, represented with colours, and modelled with LME or with PuMBA. A. Power as a function of sample size for a 0.1 difference between patients and controls. The black dashed line represents the power of the LME analysis applied to the true values of ***BP***_P_ from which the simulations were generated. B. Mean standard error of the estimated group differences across the simulated datasets. The black dashed line represents the mean standard error of the LME analysis applied to the true values of ***BP***_P_. C. Mean estimated group differences across simulated datasets, with the true difference of 0.1 shown with a dashed black line. The error bars represent the standard deviation of estimated group differences across simulations, shown only for the original measurement error.

In all cases, the power of PuMBA exceeds that of the LME model for the same degree of measurement error. In most circumstances, the power of PuMBA even exceeds that of applying LME to the “true” ***BP***_P_ values, i.e. assuming perfect quantification. PuMBA analysis yielded lower standard error (Figure 2B) as well as lower standard deviation between simulated datasets (Figure 2C and Supplementary Materials S3) of the estimated group differences between simulations for relative to LME. These imply respectively that PuMBA estimates exhibit greater precision, or certainty, of the magnitude of the group differences; and that PuMBA estimates are more consistent between simulated samples, i.e. differ less from sample to sample. Decreases in the standard error of estimates of group differences without correspondingly large decreases in the standard deviation of these estimates across samples would result in an increase in the false positive rate in the presence of no group differences. However, we see no evidence for any increase in the false positive rate when PuMBA is applied to data simulated with no group differences: in fact PuMBA exhibits a lower false positive rate on average for both tracers, and for every level of measurement error (Supplementary Materials S3).

Both LME and PuMBA exhibit a small degree of bias in the estimated group differences (Figure 2C) presumably owing to inaccuracies in the PK parameters estimated using the NLS model fitting procedure, i.e. prior to the parameters being entered into the statistical models. For [^11^C]ABP688, there was a tendency to underestimate group differences, while for [^11^C]WAY100635 there was a tendency to overestimate group differences. In all cases this bias was greater for PuMBA than for LME, suggesting that PuMBA exacerbates this bias. The bias was also more pronounced both in cases of smaller sample sizes and greater measurement error. For [^11^C]ABP688 with the original measurement error, mean estimates across the simulated datasets of the true difference of 0.1 were 0.066 and 0.080 for PuMBA and LME respectively for n=10; while for n=100 the mean estimates were 0.084 and 0.089. For [^11^C]WAY100635 with the original measurement error, the mean estimates were 0.138 and 0.104 for PuMBA and LME respectively for n=10, while for n=100 they were 0.105 and 0.101, respectively. In all cases however, the bias of the estimates was smaller than the sample-to-sample variation: the bias of the group difference estimates relative to the true value was never greater than 65% of the standard deviation of these estimates across simulated datasets (median: 41% for [^11^C]ABP688 and 23% for [^11^C]WAY100635; see Supplementary Materials S3). This implies that the combined effects of sampling error and measurement error are of greater consequence for the estimation of group differences compared to the bias in the PuMBA estimate for any given sample.

#### 3.1.1 Comparison with SiMBA

In order to directly compare the performance of PuMBA with SiMBA using the same priors, we applied both models to the same simulated datasets as described in our previous report [4]. Furthermore, in order to maximise the comparability of the outcomes, we estimated the same binding parameter, ***BP***_ND_ (as opposed to ***BP***_P_ as before) using both PuMBA and SiMBA with exactly the same priors on all shared parameters. Firstly, MCMC sampling time is much more rapid for PuMBA compared to SiMBA (Figure 3A), showing that fitting a PuMBA model is completed approximately 4000 times more quickly compared to SiMBA for the same number of iterations. We show that SiMBA outperforms PuMBA, with greater power (Figure 3B), lower standard error (Figure 3C), lower standard deviations of estimated group differences (Figure 3D), and a smaller degree of bias (Supplementary Materials S4). Lastly, while PuMBA and SiMBA both outperform univariate LME analysis of the true values of ***BP***_ND_ underlying the simulations in power, standard error or standard deviation across simulations, they do not outperform a multivariate PuMBA model fit to the true values of all four of the PK parameters.

**Fig. 3.**
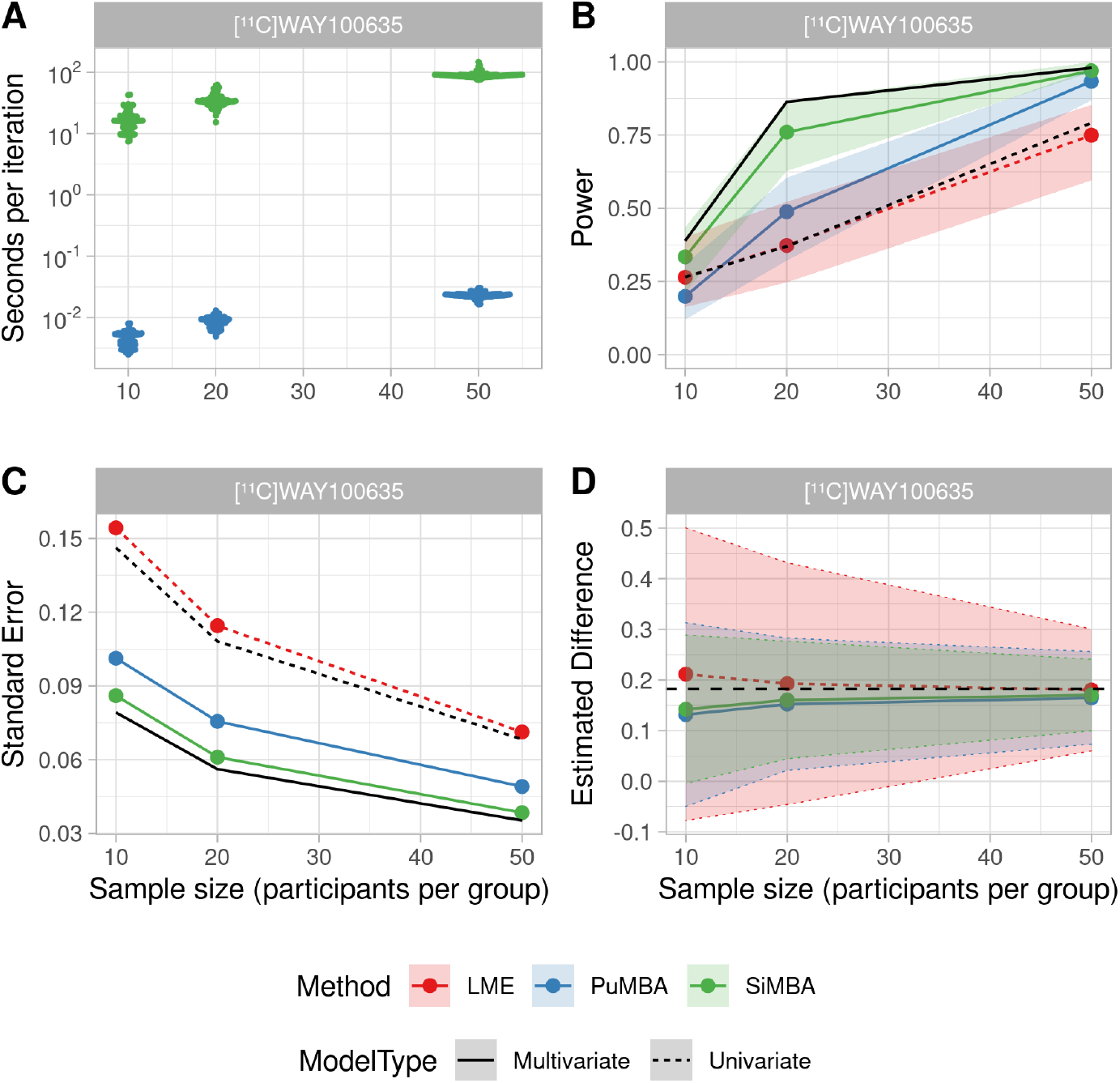
PuMBA and SiMBA were applied to the same simulated datasets in each condition, with identical priors on all shared parameters to compare their performance. Note that the measurement error sigma is equal to approximately double the original measurement error [4]. Black lines refer to the performance of these models applied to the true values underlying the simulations: univariate LME applied to the binding parameter, and multivariate PuMBA analysis applied to the true values of all the pharmacokinetic parameters. A. MCMC sampling times for PuMBA and SiMBA per iteration. B. Power as a function of sample size for a 0.182 (20%) difference between patients and controls. The shaded area represents the upper and lower bounds of the 95% confidence interval obtained using bootstrap resampling. C. Standard error of group difference estimates as a function of sample size. D. Mean estimated group differences across simulated datasets, with the true difference of 0.182 shown with a dashed black line. The shaded area represents the standard deviation of estimated group differences across simulations for each approach, with their individual boundaries emphasised with dotted lines.

### 3.2 Correlation Matrix Recovery

Since PuMBA exploits the correlations between PK parameters, it is important to consider how well these correlations are estimated and, as a result, to what extent the bias or variance in these estimates affect the quality of PuMBA inferences, especially the false negative rate. To this end, we extracted the estimated correlation coefficients from the TAC simulations described in section 3.1 to compare with the corresponding matrices that were set for the simulation. We also simulated additional uncorrelated data in which the parameters were generated using multivariate distributions but with the correlations between parameters removed, i.e. with diagonal variance-covariance matrices. Using the uncorrelated simulation parameters, we simulated both TAC data and parameter data to examine the influence of PK parameter estimation from TACs. More details are provided in Supplementary Materials S5.

For simulated uncorrelated parameter data, estimated correlations were centred around zero for all parameter pairs, demonstrating that PuMBA itself is capable of recovering the parameter intercorrelations accurately in the absence of any bias introduced during PK parameter estimation from TACs. However in simulated TAC data, the recovery of the parameter intercorrelations was reasonably poor for most pairs of PK parameters in the individual (***τ***) correlation matrices, both for the correlated and uncorrelated data, in contrast to SiMBA [4]. Together, these results imply that the poor recovery of the true parameter correlations is primarily due to bias in the estimation of the PK parameters from TACs using NLS.

Despite the poor recovery of parameter intercorrelations, application of PuMBA to uncorrelated data resulted in higher mean standard error and standard deviation of group difference estimates, and reductions in statistical power relative to the original correlated data. In the correlated simulations, estimated correlation coefficients were closer to the true simulated values for [^11^C]ABP688 than for [^11^C]WAY100635, which may explain the greater improvements performance observed with PuMBA relative to LME with [^11^C]ABP688. Together, these results suggest that the power and precision of PuMBA estimates are influenced by the true underlying parameter intercorrelations, and are likely improved when these intercorrelations are more accurately estimated.

PuMBA therefore exploits parameter intercorrelations to improve its inferences, yet these correlations tend to be estimated relatively poorly. We were concerned that this might imply that, in the absence of true correlations, that the artefactual correlations arising from parameter estimation bias might yield a higher risk of false positive conclusions. However, for both [^11^C]ABP688 and [^11^C]WAY100635 we observed no apparent increase in the false positive rate in either of the uncorrelated datasets relative to either PuMBA applied to the correlated datasets or to LME (Supplementary Materials S5).

### 3.3 Parameter Simulations

In order to test whether PuMBA can also be applied for other kinetic models which cannot yet be modelled using SiMBA, we performed additional parameter-only simulations using the 1TC and SRTM, as described in section 2.4.2. The results of these simulations are shown in Figure 4, in which we observe increases in power for linear mixed effects (LME) modelling relative to *t*-tests, and improved power for PuMBA relative to LME. We observe larger improvements in power for SRTM, while improvements in power for the 1TC were more modest. In all cases, these improvements in power were not associated with any increase in false positive rate (Supplementary Materials S6).

**Fig. 4.**
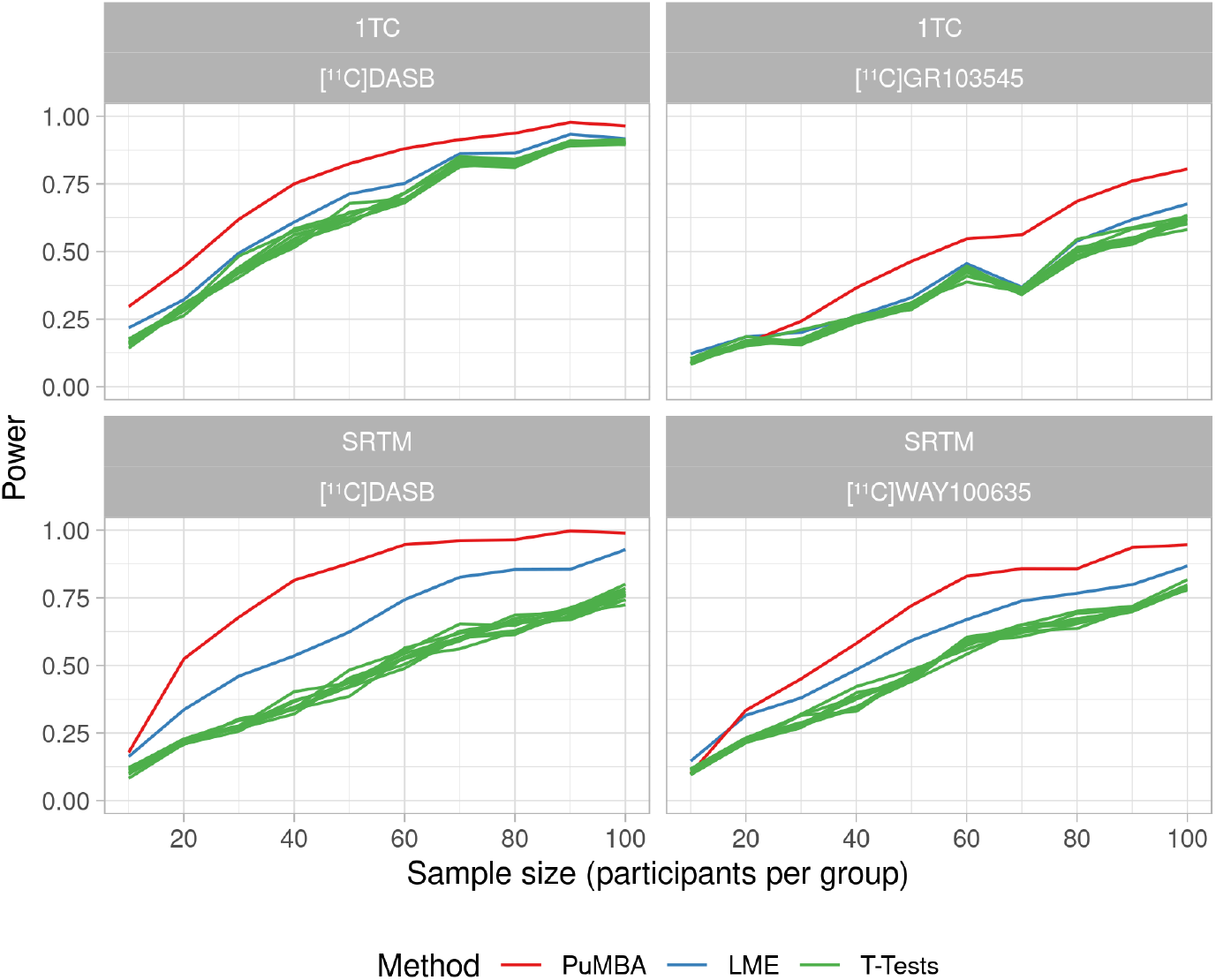
Power as a function of sample size for a 0.1 difference between patients and controls. The models are the one-tissue compartment model (1TC) and the simplified reference tissue model (SRTM).

When examining the estimated group differences, we observed lower standard error as well as lower standard deviation of the estimated group differences between simulated datasets for LME relative to *t*-tests, as well as for PuMBA relative to LME. While we observed no bias in the mean estimated group differences for *t*-tests or LME, PuMBA showed slightly biased estimates in all cases, with more bias in smaller sample sizes (Supplementary Materials S6).

## 4 Discussion

In this study, we show that a multivariate statistical analysis of all estimated PET pharmacokinetic parameters using PuMBA yields significant inferential advantages over univariate analysis of the parameter of interest performed without considering the other parameters. We also show that PuMBA can be fruitfully applied even when there is a substantial degree of measurement error. As would be expected, because PuMBA is applied to parameter estimates, it cannot outperform a similar analysis which is conducted on the full TAC data in which quantification and statistical analysis are both able to benefit from the hierarchical multivariate framework, i.e. SiMBA. However, PuMBA models can be fitted in minutes, in contrast to days required by SiMBA. Furthermore, PuMBA can be directly applied to data from more PET pharmacokinetic models as a substitute for conventional statistical analysis approaches, while SiMBA requires the user to incorporate the pharmacokinetic model itself into the Bayesian model, which can be challenging.

While PuMBA can more easily be applied to a wider range of pharmacokinetic models than SiMBA, this does not necessarily imply that it can be applied when the identifiability of PK parameters for a given model and tracer is poor. For instance, the identifiability of the 2TC is reasonably good for [^11^C]WAY100635 and [^11^C]ABP688, in contrast with [^11^C]GR103545 for which the 2TC is rather poorly identified [33]. On the other hand, SiMBA stabilises the fitting of the PK model itself using the hierarchical multifactor framework, and therefore has the additional advantage of improving model identifiability: this is not a property of PuMBA. Hence, while SiMBA is more generalisable in the sense that it makes the application of the more complex models possible for a wider variety of PET tracers for which they are otherwise insufficiently identifiable, PuMBA is more generalisable in the sense that it can, without any substantive modifications, be applied to data originating from a wider assortment of PK models, including reference tissue models – provided that the parameters are sufficiently identifiable.

Since PuMBA, in contrast to SiMBA, cannot improve the estimation of the PK parameters themselves from the NLS model, its function is only to take advantage of information shared between individuals, regions and parameters. Both LME and PuMBA make use of partial pooling, and both make use of the same predictors for the binding parameter. Thus, the only difference between the performance of these models is due to the multivariate partial pooling strategy applied in PuMBA, in contrast to the univariate partial pooling applied in LME. Moreoever, by exploiting the relationships between PK parameters, PuMBA exhibits greater precision and power compared to a univariate LME analysis of even the *true* binding parameters from which the simulations were generated, i.e. as if binding could be estimated perfectly. This can be described with an analogy. Consider that we wish to compare binding potential of two groups of individuals. We have the option of either measuring all of our participants in a noisy PET system and modelling their resulting TACs — or of being informed of each individual’s true binding potential value by an omniscient oracle. Counter-intuitively, the present results suggest that we ought to shun the oracle, and perform the PET study as planned using PuMBA, or SiMBA, for the statistical analysis. However, if the oracle could be persuaded to provide us with the true values of *all* of the PK parameters, then of course it would be advantageous to accept the oracle’s help. This corresponds to the comparisons of SiMBA and PuMBA with the solid and dashed black lines in Figures 3B–C. In other words, by discarding the otherwise irrelevant PK parameters during univariate statistical analysis of the parameter representing target binding, we effectively remove valuable context for our model. We are therefore guilty of floccinaucinihilipilification when it comes to the additional estimated PK parameters from our models.

It cannot be assumed that a multivariate analysis will improve the power of any given statistical comparison. With reference to Figure 1, it is clear that with sufficiently strong correlations between parameters, and sufficiently precise estimates of the parameters and their correlations with one another, the variance of the conditional distribution of an estimated parameter can be reduced to below that of the marginal distribution of the true parameter. However, in the presence of high uncertainty, small samples, or low correlation between variables, this potential gain is likely to be reduced: we see some evidence of this with *n* = 10 in Figures 2A and 3B. We also see that PuMBA yielded greater improvements in precision and statistical power when the correlation matrices were more accurately estimated in [^11^C]ABP688 compared to [^11^C]WAY100635.

In all applications, PuMBA exhibited bias in its group difference estimates, particularly for smaller sample sizes and with larger measurement error, for instance in Figure 2C. We find that this bias was never greater than 65% of the standard deviation of sample-to-sample estimate variability, which implies that even in the worst case scenario, the estimated differences for [^11^C]ABP688 will still be greater than the true value approximately 25% of the time given infinite replications of the same study. Although the magnitude of this bias is not large, it is still important to understand its source. One factor is that PuMBA makes use of a regularising prior for the estimation of group differences, which makes the model skeptical of large differences between groups, effectively shrinking the estimated group differences towards zero. However, for [^11^C]WAY100635, we observe that the bias is positive, which is incompatible with shrinkage being solely responsible for the observed bias. This corresponds with the observation that the LME group differences are also biased in the same direction as the PuMBA estimates in the TAC simulations, but to a smaller degree in each case. This suggests that the bias is also partially explained by identifiability issues in the PK parameter estimation from the TAC data using NLS, and that bias in the estimation of the binding parameter is also accompanied by bias in the estimation of the other model parameters in a systematic fashion. This corresponds with the results of the uncorrelated data simulations, in which correlations were partially induced between parameters by the PK parameter estimation from TACs. It is noteworthy however that PuMBA exacerbates this bias, presumably owing to its estimation of the correlations between the PK parameter estimates with potentially induced correlations. Finally, for 1TC parameter simulations of [^11^C]DASB, there is positive bias in small sample sizes, which cannot be explained by either of the above reasons. This suggests that the identifiability of the PuMBA model itself may also play a role in the observed bias, particularly in small sample sizes. For these reasons, the precise magnitude of PuMBA estimates should be interpreted with some degree of care particularly in applications for which the identifiability of the individual parameters is poor, or in small sample sizes.

While PuMBA is much faster to estimate and easier to implement compared to SiMBA, it does still require additional effort from the analyst compared to conventional approaches. Firstly, we would advise paying more attention to the estimated parameters generated by the NLS procedures prior to analysis: even when estimates of binding potential, for example, may appear reasonable, estimates for some of the other parameters could be problematic. For this reason, we made use of reasonably conservative upper and lower limits when fitting the NLS models, coupled with estimating the fit with multiple starting points [34] in order to minimise the chance of our model yielding parameter estimates obtained from a local minimum. Another additional requirement of PuMBA compared to conventional approaches is that, because all parameters are included in the PuMBA model, the analyst must define linear model specifications for each of them. Age, for instance, might affect blood delivery, or the kinetics of the radioligand in a reference tissue, even if it does not affect the binding potential for a particular target. Furthermore, because PuMBA is a Bayesian model, prior distributions need to be defined not only for all covariates, but for all parameters in the model. Using PuMBA therefore requires more care and consideration than conventional approaches, and in order to be applied most effectively, we recommend collaboration between researchers with domain expertise and technical expertise. This allows for the incorporation what is already known about the relevant clinical and biological constraints into the specification and priors of the model for a given research question.

One interesting observation which emerges from the fitting of all of the datasets is the fact that inter-individual variation in blood delivery (***K***_1_ and **R**_1_) and binding (***V***_t_, ***BP***_P_ and ***BP***_ND_) were positively correlated with one another across all models and tracers, using both PuMBA and SiMBA (Supplementary Materials S2). Importantly, these correlations refer to the correlation of partially pooled estimates (i.e. random effects) of the PK parameters at the individual level, and not to the parameters estimated for each individual TAC. While PuMBA may be more sensitive to issues of poor identifiability when fitting the PK model, SiMBA ought to be more robust to this possibility. The exact meaning of this observation is beyond the scope of the current investigation, and perhaps cannot be understood using pharmacokinetic modelling alone. However, this raises questions about how independent the parameters of these models are, or ought to be between individuals; as well as whether this has implications for how these parameters can be understood.

In conclusion, PuMBA allows researchers to make more precise and powerful statistical inferences, and thereby extract more information from a given dataset without needing to collect any additional information. PuMBA takes in the order of minutes to fit on a single computer core and can more easily be applied to a wider range of pharmacokinetic models than SiMBA. PuMBA may therefore serve as a convenient intermediate substitute for a full SiMBA analysis, and a useful tool for statistical analysis of PET data in general. Potential avenues for future research include examining the similarity of estimated parameter intercorrelations between different datasets collected by different groups, assessing the factors which contribute to the differential performance of PuMBA in different settings, as well as whether PuMBA could be applied simultaneously to data collected using multiple radioligands in the same individuals to estimate and take advantage of similarities in blood delivery or in target protein levels within individuals.

## 5 Acknowledgements

The work reported here has been partially supported by US NIH grants 5 P50 MH090964 and 5 R01EB024526, Hjärnfonden (PS2020-0016) and Vetenskapsrådet (2020-06356). We also thank the MIND group, and Francesca Zanderigo in particular, for their thought-provoking discussions and input, as well as provision of data.

