## Supplementary Materials for "Multivariate Analysis of PET Pharmacokinetic Parameters"

**Supplementary Materials S3 : TAC Simulation Additional Figures** **17**

**Supplementary Materials S4 : Comparison with SiMBA Additional Figures** **20**

**Supplementary Materials S5 : Correlated and Uncorrelated Data** **22**

**Supplementary Materials S6 : Parameter Simulation Additional Figures** **27**

### Supplementary Materials S1 : Prior Specification

#### Global Intercepts

Below are the priors defined for the global intercepts. Note that all priors are defined over the natural logarithms of the parameters.

##### Two-Tissue Compartment Model

**[$^{11}$C]WAY100635**

$$\begin{aligned} \alpha_{\mathrm{K_1}} &\sim \mathrm{Normal}(-2.5\ ,\ 0.25) \\ \alpha_{\mathrm{V_{ND}}} &\sim \mathrm{Normal}(-1.0\ ,\ 0.25) \\ \alpha_{\mathrm{BP_P}} &\sim \mathrm{Normal}(\ \ 1.0\ ,\ 0.25) \\ \alpha_{\mathrm{k_4}} &\sim \mathrm{Normal}(-4.0\ ,\ 0.25) \end{aligned}$$

The priors for $\alpha_{\mathrm{K_1}}$ and $\alpha_{\mathrm{BP_P}}$ are defined for the dorsolateral prefrontal cortex as the reference level of the dummy variable.

**[$^{11}$C]ABP688**

$$\begin{aligned} \alpha_{\mathrm{K_1}} &\sim \mathrm{Normal}(-2.0\ ,\ 0.25) \\ \alpha_{\mathrm{V_{ND}}} &\sim \mathrm{Normal}(-0.5\ ,\ 0.5) \\ \alpha_{\mathrm{BP_P}} &\sim \mathrm{Normal}(\ \ 0.5\ ,\ 0.5) \\ \alpha_{\mathrm{k_4}} &\sim \mathrm{Normal}(-2.5\ ,\ 0.5) \end{aligned}$$

The priors for $\alpha_{\mathrm{K_1}}$ and $\alpha_{\mathrm{BP_P}}$ are defined for the dorsolateral prefrontal cortex as the reference level of the dummy variable.

##### One-Tissue Compartment Model

**[$^{11}$C]DASB**

$$\begin{aligned} \alpha_{\mathrm{K_1}} &\sim \mathrm{Normal}(-1\ ,\ 0.25) \\ \alpha_{\mathrm{V_T}} &\sim \mathrm{Normal}(2\ ,\ 0.25) \end{aligned}$$

The priors for $\alpha_{\mathrm{K_1}}$ and $\alpha_{\mathrm{V_T}}$ are defined for the anterior cingulate cortex as the reference level of the dummy variable.

**[$^{11}$C]GR103545**

$$\begin{aligned} \alpha_{\mathrm{K_1}} &\sim \mathrm{Normal}(-2.0\ ,\ 0.25) \\ \alpha_{\mathrm{V_T}} &\sim \mathrm{Normal}(3\ ,\ 0.5) \end{aligned}$$

The priors for $\alpha_{\mathrm{K_1}}$ and $\alpha_{\mathrm{V_T}}$ are defined for the dorsolateral prefrontal cortex as the reference level of the dummy variable.

##### Simplified Reference Tissue Model

###### [$^{11}$C]DASB

$$\begin{aligned}
\alpha_{\mathrm{R_1}} &\sim \mathrm{Normal}(0\,,\,0.25)\\
\alpha_{\mathrm{k'_2}} &\sim \mathrm{Normal}(-3\,,\,0.25)\\
\alpha_{\mathrm{BP_{ND}}} &\sim \mathrm{Normal}(-1\,,\,0.25)
\end{aligned}$$

The priors for $\alpha_{\mathrm{R_1}}$ and $\alpha_{\mathrm{BP_{ND}}}$ are defined for the anterior cingulate cortex as the reference level of the dummy variable.

###### [$^{11}$C]WAY100635

$$\begin{aligned}
\alpha_{\mathrm{R_1}} &\sim \mathrm{Normal}(0\,,\,0.25)\\
\alpha_{\mathrm{k'_2}} &\sim \mathrm{Normal}(-2.3\,,\,0.25)\\
\alpha_{\mathrm{BP_{ND}}} &\sim \mathrm{Normal}(1.5\,,\,0.25)
\end{aligned}$$

The priors for $\alpha_{\mathrm{R_1}}$ and $\alpha_{\mathrm{BP_{ND}}}$ are defined for the dorsolateral prefrontal cortex as the reference level of the dummy variable.

#### Individual deviations

Differences between individuals were defined by specifying the primary pharmacokinetic parameters in one variance-covariance matrix.

##### Two-Tissue Compartment Model

For both [$^{11}$C]WAY100635 and [$^{11}$C]ABP688, we used the same priors.

$$\begin{bmatrix}\tau_{\mathrm{K_1}}\\ \tau_{\mathrm{V_{ND}}}\\ \tau_{\mathrm{BP_{ND}}}\\ \tau_{\mathrm{k_4}}\end{bmatrix} \sim \mathrm{MVNormal}\left(\begin{bmatrix}0\\0\\0\\0\end{bmatrix}, \boldsymbol{\Sigma}_{\mathrm{Subject}}\right)$$

$$\Sigma_{\mathrm{Subject}} = \begin{bmatrix}\sigma_{\mathrm{K_1}} & 0 & 0\\ 0 & \ddots & 0\\ 0 & 0 & \sigma_{\mathrm{k_4}}\end{bmatrix} \mathbf{R}_{\mathrm{Subject}} \begin{bmatrix}\sigma_{\mathrm{K_1}} & 0 & 0\\ 0 & \ddots & 0\\ 0 & 0 & \sigma_{\mathrm{k_4}}\end{bmatrix}$$

$$\begin{aligned}
\sigma_{\mathrm{K_1}} &\sim \text{Half-Normal}(0\,,\,0.3)\\
\sigma_{\mathrm{V_{ND}}} &\sim \text{Half-Normal}(0\,,\,0.3)\\
\sigma_{\mathrm{BP_{ND}}} &\sim \text{Half-Normal}(0\,,\,0.3)\\
\sigma_{\mathrm{k_4}} &\sim \text{Half-Normal}(0\,,\,0.3)\\
\mathbf{R}_{\mathrm{Subject}} &\sim \mathrm{LKJ}(1)
\end{aligned}$$

##### One-Tissue Compartment Model

For both [$^{11}$C]DASB and [$^{11}$C]GR103545, we used the same priors.

$$
\begin{aligned}
\begin{bmatrix} \tau_{\mathrm{K_1}} \\ \tau_{\mathrm{V_T}} \end{bmatrix} &\sim \mathrm{MVNormal}\left( \begin{bmatrix} 0 \\ 0 \end{bmatrix}, \mathbf{\Sigma}_{\mathrm{Subject}} \right) \\
\Sigma_{\mathrm{Subject}} &= \begin{bmatrix} \sigma_{\mathrm{K_1}} & 0 \\ 0 & \sigma_{\mathrm{V_T}} \end{bmatrix} \mathbf{R}_{\mathrm{Subject}} \begin{bmatrix} \sigma_{\mathrm{K_1}} & 0 \\ 0 & \sigma_{\mathrm{V_T}} \end{bmatrix} \\
\\
\sigma_{\mathrm{K_1}} &\sim \text{Half-Normal}(0\,,\ 0.3) \\
\sigma_{\mathrm{V_T}} &\sim \text{Half-Normal}(0\,,\ 0.3) \\
\mathbf{R}_{\mathrm{Subject}} &\sim \mathrm{LKJ}(1)
\end{aligned}
$$

##### Simplified Reference Tissue Model

For both [$^{11}$C]DASB and [$^{11}$C]WAY100635, we used the same priors.

$$
\begin{aligned}
\begin{bmatrix} \tau_{\mathrm{R_1}} \\ \tau_{\mathrm{k_2'}} \\ \tau_{\mathrm{BP_{ND}}} \end{bmatrix} &\sim \mathrm{MVNormal}\left( \begin{bmatrix} 0 \\ 0 \\ 0 \end{bmatrix}, \mathbf{\Sigma}_{\mathrm{Subject}} \right) \\
\Sigma_{\mathrm{Subject}} &= \begin{bmatrix} \sigma_{\mathrm{R_1}} & 0 & 0 \\ 0 & \sigma_{\mathrm{k_2'}} & 0 \\ 0 & 0 & \sigma_{\mathrm{BP_{ND}}} \end{bmatrix} \mathbf{R}_{\mathrm{Subject}} \begin{bmatrix} \sigma_{\mathrm{R_1}} & 0 & 0 \\ 0 & \sigma_{\mathrm{k_2'}} & 0 \\ 0 & 0 & \sigma_{\mathrm{BP_{ND}}} \end{bmatrix} \\
\\
\sigma_{\mathrm{R_1}} &\sim \text{Half-Normal}(0\,,\ 0.3) \\
\sigma_{\mathrm{k_2'}} &\sim \text{Half-Normal}(0\,,\ 0.3) \\
\sigma_{\mathrm{BP_{ND}}} &\sim \text{Half-Normal}(0\,,\ 0.3) \\
\mathbf{R}_{\mathrm{Subject}} &\sim \mathrm{LKJ}(1)
\end{aligned}
$$

#### Regional deviations

For $\log\mathrm{BP_{ND}}$ and $\log\mathrm{K_1}$, regional differences were defined as unpooled effects using a dummy (indicator) variable defined with reference to the dorsolateral prefrontal cortex. For simplicity, all regional differences (with the exception of [$^{11}$C]DASB) were defined as zero-centred regularising priors with the same SD.

$$
\begin{aligned}
\upsilon_{\mathrm{j,K_1}} &\sim \mathrm{Normal}(0\,,\ 0.3) \\
\upsilon_{\mathrm{j,BP_{ND}}} &\sim \mathrm{Normal}(0\,,\ 0.3)
\end{aligned}
$$

For [$^{11}$C]DASB, the same standard deviations were used, but with altered means for regions whose mean binding values were very different from the other regions. For SRTM, the mean values for $$ were defined as follows:

- Insula: 0.5

- Amygdala, dorsal putamen, thalamus, ventral striatum: 1
- Midbrain: 1.5

For the 1TC, the means were defined as follows:

- Dorsal putamen, ventral striatum: 0.5
- Midbrain: 1

For the remaining parameters, regional differences were defined as pooled variables, arising from a common distribution

$$\begin{aligned}
\begin{bmatrix} v \\ \vdots \end{bmatrix} &\sim \text{MVNormal}\left(\begin{bmatrix} 0 \\ \vdots \end{bmatrix}, \mathbf{\Sigma}_{\text{Region}}\right) \\
\Sigma_{\text{Region}} &= \begin{bmatrix} \sigma \\ \vdots \end{bmatrix} \mathbf{R}_{\text{Region}} \begin{bmatrix} \sigma \\ \vdots \end{bmatrix} \\
\sigma &\sim \text{Half-Normal}(0\ ,\ 0.1) \\
\mathbf{R}_{\text{Region}} &\sim \text{LKJ}(2)
\end{aligned}$$

#### Residuals

##### Two-Tissue Compartment Model

For both $[^{11}C]$WAY100635 and $[^{11}C]$ABP688, we used the same priors.

$$\begin{aligned}
\begin{bmatrix} \epsilon_{\text{K}_1} \\ \epsilon_{\text{V}_{\text{ND}}} \\ \epsilon_{\text{BP}_{\text{ND}}} \\ \epsilon_{\text{k}_4} \end{bmatrix} &\sim \text{MVNormal}\left(\begin{bmatrix} 0 \\ 0 \\ 0 \\ 0 \end{bmatrix}, \mathbf{\Sigma}_{\text{Subject}}\right) \\
\Sigma_{\text{Subject}} &= \begin{bmatrix} \sigma_{\text{K}_1} & 0 & 0 \\ 0 & \ddots & 0 \\ 0 & 0 & \sigma_{\text{k}_4} \end{bmatrix} \mathbf{R}_{\text{Subject}} \begin{bmatrix} \sigma_{\text{K}_1} & 0 & 0 \\ 0 & \ddots & 0 \\ 0 & 0 & \sigma_{\text{k}_4} \end{bmatrix} \\
\sigma_{\text{K}_1} &\sim \text{Half-Normal}(0\ ,\ 0.1) \\
\sigma_{\text{V}_{\text{ND}}} &\sim \text{Half-Normal}(0\ ,\ 0.1) \\
\sigma_{\text{BP}_{\text{ND}}} &\sim \text{Half-Normal}(0\ ,\ 0.1) \\
\sigma_{\text{k}_4} &\sim \text{Half-Normal}(0\ ,\ 0.1) \\
\mathbf{R}_{\text{Subject}} &\sim \text{LKJ}(2)
\end{aligned}$$

##### One-Tissue Compartment Model

For both $[^{11}C]$DASB and $[^{11}C]$GR103545, we used the same priors.

$$
\begin{aligned}
\begin{bmatrix} \epsilon_{\mathrm{K_1}} \\ \epsilon_{\mathrm{V_T}} \end{bmatrix} &\sim \mathrm{MVNormal}\left( \begin{bmatrix} 0 \\ 0 \end{bmatrix}, \boldsymbol{\Sigma}_{\mathrm{Subject}} \right) \\
\Sigma_{\mathrm{Subject}} &= \begin{bmatrix} \sigma_{\mathrm{K_1}} & 0 \\ 0 & \sigma_{\mathrm{V_T}} \end{bmatrix} \mathbf{R}_{\mathrm{Subject}} \begin{bmatrix} \sigma_{\mathrm{K_1}} & 0 \\ 0 & \sigma_{\mathrm{V_T}} \end{bmatrix} \\
\sigma_{\mathrm{K_1}} &\sim \text{Half-Normal}(0\ ,\ 0.1) \\
\sigma_{\mathrm{V_T}} &\sim \text{Half-Normal}(0\ ,\ 0.1) \\
\mathbf{R}_{\mathrm{Subject}} &\sim \mathrm{LKJ}(2)
\end{aligned}
$$

##### Simplified Reference Tissue Model

For both [$^{11}$C]DASB and [$^{11}$C]WAY100635, we used the same priors.

$$
\begin{aligned}
\begin{bmatrix} \epsilon_{\mathrm{R_1}} \\ \epsilon_{\mathrm{k_2'}} \\ \epsilon_{\mathrm{BP_{ND}}} \end{bmatrix} &\sim \mathrm{MVNormal}\left( \begin{bmatrix} 0 \\ 0 \\ 0 \end{bmatrix}, \boldsymbol{\Sigma}_{\mathrm{Subject}} \right) \\
\Sigma_{\mathrm{Subject}} &= \begin{bmatrix} \sigma_{\mathrm{R_1}} & 0 & 0 \\ 0 & \sigma_{\mathrm{k_2'}} & 0 \\ 0 & 0 & \sigma_{\mathrm{BP_{ND}}} \end{bmatrix} \mathbf{R}_{\mathrm{Subject}} \begin{bmatrix} \sigma_{\mathrm{R_1}} & 0 & 0 \\ 0 & \sigma_{\mathrm{k_2'}} & 0 \\ 0 & 0 & \sigma_{\mathrm{BP_{ND}}} \end{bmatrix} \\
\sigma_{\mathrm{R_1}} &\sim \text{Half-Normal}(0\ ,\ 0.1) \\
\sigma_{\mathrm{k_2'}} &\sim \text{Half-Normal}(0\ ,\ 0.1) \\
\sigma_{\mathrm{BP_{ND}}} &\sim \text{Half-Normal}(0\ ,\ 0.1) \\
\mathbf{R}_{\mathrm{Subject}} &\sim \mathrm{LKJ}(2)
\end{aligned}
$$

#### Covariates

For the simulations, there was only one additional covariate for group effects on the binding parameter. For this, we used a zero-centred regularising prior. This prior conservatively assumes that no difference is most likely, and assigns 95% of its probability between differences of -48% and 48% differences between groups.

$$\beta_{\mathrm{Group}} \sim \mathrm{Normal}(\ 0\ ,\ 0.2\ )$$

### Supplementary Materials S2 : Simulation Parameters

#### Global Intercepts

##### Two-Tissue Compartment Model

[$^{11}$C]WAY100635

| Parameter | Mean |
|---|---|
| logK1 | -2.33 |
| logBPnd | 1.75 |
| logVnd | -0.62 |
| logk4 | -3.84 |
| logvB | -3.63 |
| logsigma | -4.96 |

[$^{11}$C]ABP688

| Parameter | Mean |
|---|---|
| logK1 | -1.95 |
| logBPnd | 0.86 |
| logVnd | -0.33 |
| logk4 | -2.57 |
| logvB | -3.26 |
| logsigma | -5.08 |

##### One-Tissue Compartment Model

[$^{11}$C]DASB

| Parameter | Mean |
|---|---|
| logK1 | -0.76 |
| logVt | 2.73 |

[$^{11}$C]GR103545

| Parameter | Mean |
|---|---|
| logK1 | -1.87 |
| logVt | 3.05 |

##### Simplified Reference Tissue Model

[$^{11}$C]DASB

| Parameter | Mean |
|---|---|
| logR1 | -0.08 |
| logk2prime | -3.06 |
| logBPnd | -0.86 |

$[^{11}C]$WAY100635

| Parameter | Mean |
|---|---|
| logR1 | 0.23 |
| logk2prime | -2.52 |
| logBPnd | 1.60 |

#### Regional Deviations

##### Two-Tissue Compartment Model

[$^{11}$C]WAY100635

| Region | logK1 | logBPnd | logVnd | logk4 | logvB | logsigma |
|---|---|---|---|---|---|---|
| DLPFC | 0.00 | 0.00 | -0.09 | 0.17 | -0.07 | -0.30 |
| MPFC | -0.01 | 0.03 | -0.06 | 0.14 | 0.06 | -0.23 |
| Hippocampus | -0.22 | 0.18 | 0.29 | -0.22 | -0.07 | 0.00 |
| Amygdala | -0.33 | 0.10 | 0.01 | -0.08 | 0.00 | 0.12 |
| Parahippocampus | -0.27 | 0.00 | 0.36 | 0.08 | 0.02 | -0.10 |
| Insula | -0.04 | 0.36 | 0.01 | 0.01 | 0.17 | -0.06 |
| ACC | -0.04 | 0.15 | -0.05 | 0.09 | 0.15 | -0.11 |
| PCC | -0.02 | -0.01 | -0.10 | 0.12 | 0.13 | -0.01 |
| DRN | -0.30 | -0.03 | -0.31 | -0.30 | -0.40 | 0.69 |

[$^{11}$C]ABP688

| Region | logK1 | logBPnd | logVnd | logk4 | logvB | logsigma |
|---|---|---|---|---|---|---|
| DLPFC | 0.00 | 0.00 | -0.05 | -0.01 | 0.01 | -0.32 |
| Amygdala | -0.26 | -0.04 | -0.04 | -0.06 | -0.04 | 0.16 |
| Hippocampus | -0.26 | 0.01 | -0.12 | -0.01 | -0.08 | -0.18 |
| MPFC | 0.01 | 0.00 | 0.02 | 0.05 | 0.08 | -0.15 |
| Dorsal Putamen | 0.21 | -0.13 | 0.09 | -0.02 | -0.04 | 0.04 |
| Ventral Striatum | 0.06 | 0.02 | 0.07 | -0.02 | -0.08 | 0.36 |
| Insula | 0.05 | 0.18 | -0.01 | 0.00 | 0.05 | -0.05 |
| PCC | 0.09 | 0.12 | -0.05 | 0.01 | 0.05 | 0.10 |
| ACC | 0.08 | 0.11 | 0.08 | 0.06 | 0.05 | 0.02 |

##### One-Tissue Compartment Model

[$^{11}$C]DASB

| Region | logK1 | logVt |
|---|---|---|
| ACC | 0.00 | 0.00 |
| Dorsal Putamen | 0.11 | 0.49 |
| Ventral Striatum | 0.01 | 0.63 |
| Amygdala | -0.24 | 0.51 |
| Thalamus | 0.12 | 0.43 |
| Hippocampus | -0.21 | 0.04 |
| PCC | 0.04 | 0.00 |
| Insula | 0.01 | 0.17 |
| Midbrain | -0.19 | 0.84 |

[$^{11}$C]GR103545

| Region | logK1 | logVt |
|---|---|---|
| DLPFC | 0.00 | 0.00 |
| Amygdala | -0.51 | 0.15 |
| Dorsal Putamen | -0.09 | -0.29 |
| Hippocampus | -0.33 | -0.65 |
| Insula | -0.19 | 0.20 |
| MPFC | -0.10 | 0.08 |
| Parahippocampus | -0.48 | -0.13 |
| DRN | -0.35 | -0.88 |
| Ventral Striatum | -0.20 | 0.16 |

##### Simplified Reference Tissue Model

[$^{11}$C]DASB

| Region | logR1 | logk2prime | logBPnd |
|---|---|---|---|
| ACC | 0.00 | -0.16 | 0.00 |
| Dorsal Putamen | 0.05 | 0.21 | 1.06 |
| Ventral Striatum | -0.04 | 0.17 | 1.28 |
| Amygdala | -0.26 | 0.07 | 1.11 |
| Thalamus | 0.08 | 0.13 | 0.97 |
| Hippocampus | -0.17 | -0.29 | 0.26 |
| PCC | 0.05 | -0.24 | 0.05 |
| Insula | -0.01 | 0.01 | 0.45 |
| Midbrain | -0.21 | 0.07 | 1.63 |

[$^{11}$C]WAY100635

| Region | logR1 | logk2prime | logBPnd |
|---|---|---|---|
| DLPFC | 0.00 | 0.00 | 0.00 |
| MPFC | 0.02 | -0.01 | 0.07 |
| Hippocampus | -0.11 | 0.10 | 0.47 |
| Amygdala | -0.19 | 0.02 | 0.14 |
| Parahippocampus | -0.13 | 0.20 | 0.41 |
| Insula | 0.02 | 0.07 | 0.42 |
| ACC | 0.02 | 0.00 | 0.19 |
| PCC | 0.02 | -0.09 | 0.01 |
| DRN | -0.27 | -0.34 | -0.24 |

#### Covariance Matrices

##### Two-Tissue Compartment Model

[$^{11}$C]WAY100635

#### 2TC WAY

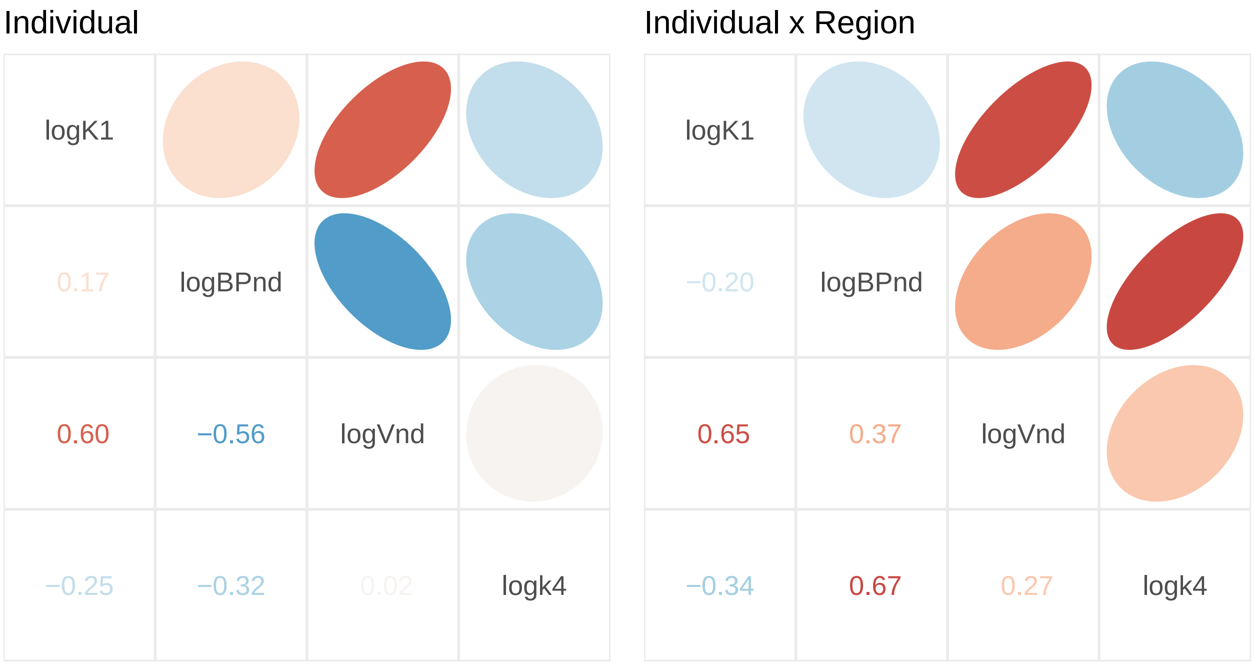

Standard Deviations

| Grouping | logK1 | logBPnd | logVnd | logk4 |
|---|---|---|---|---|
| Individual | 0.271 | 0.329 | 0.392 | 0.119 |
| Individual x Region | 0.053 | 0.037 | 0.054 | 0.016 |

$[^{11}C]$ABP688

#### 2TC ABP

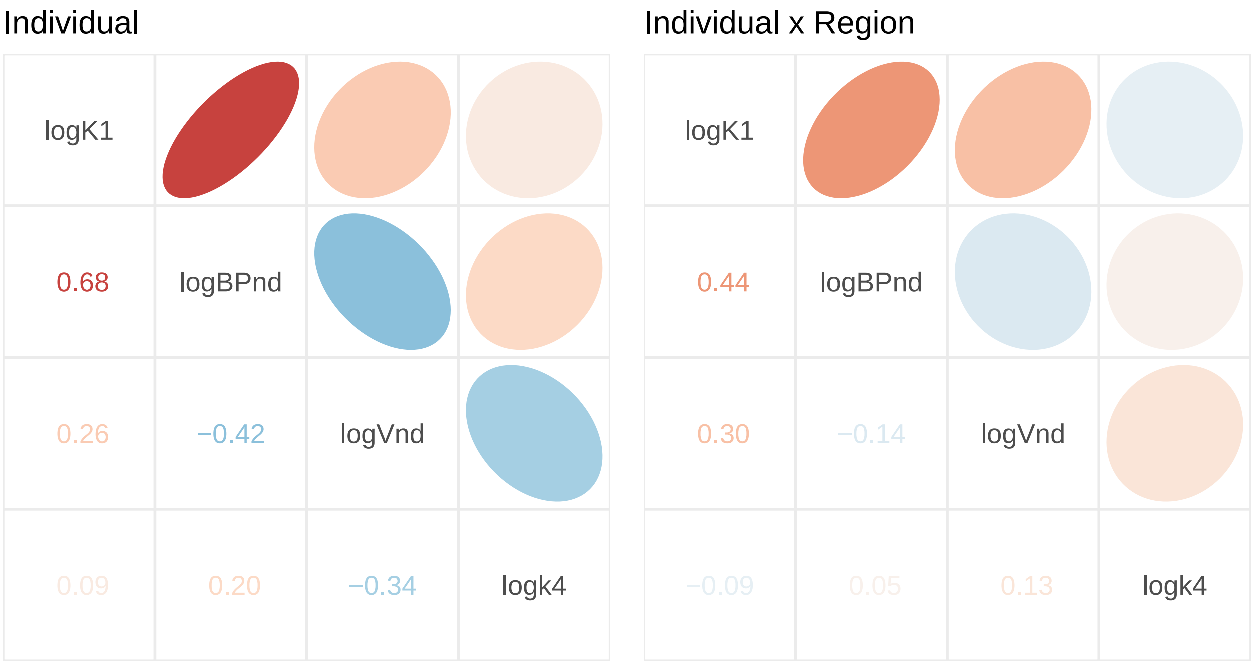

Standard Deviations

| Grouping | logK1 | logBPnd | logVnd | logk4 |
|---|---|---|---|---|
| Individual | 0.284 | 0.344 | 0.235 | 0.258 |
| Individual x Region | 0.038 | 0.035 | 0.032 | 0.032 |

**One-Tissue Compartment Model**

[$^{11}$C]DASB

#### 1TC DASB

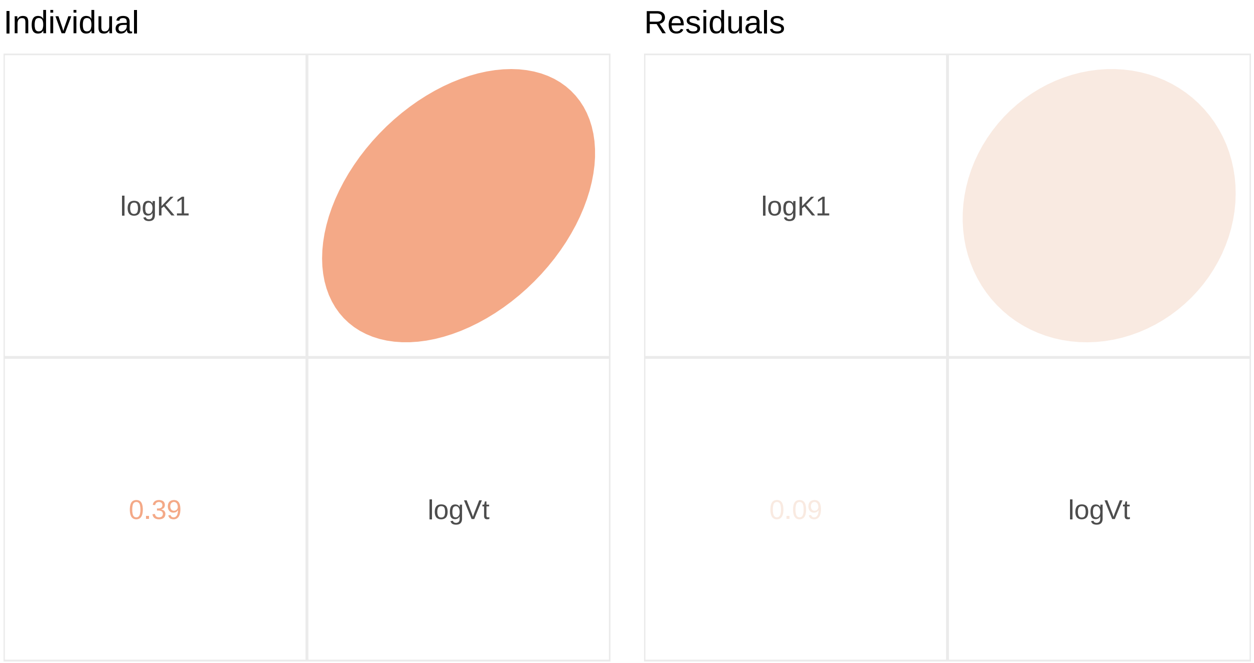

Standard Deviations

| Grouping | logK1 | logVt |
|---|---|---|
| Individual | 0.208 | 0.203 |
| Residual | 0.064 | 0.066 |

[$^{11}$C]GR103545

## 1TC GR

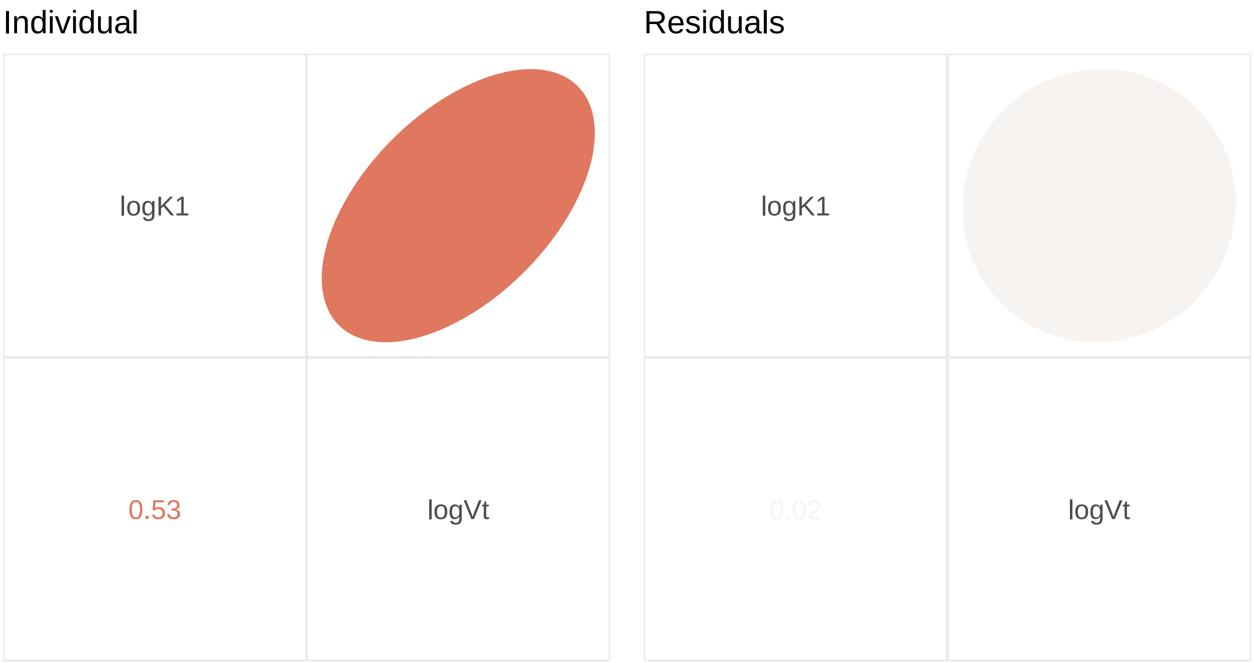

Standard Deviations

| Grouping | logK1 | logVt |
|---|---|---|
| Individual | 0.266 | 0.312 |
| Residual | 0.041 | 0.103 |

##### Simplified Reference Tissue Model

$[^{11}C]$DASB

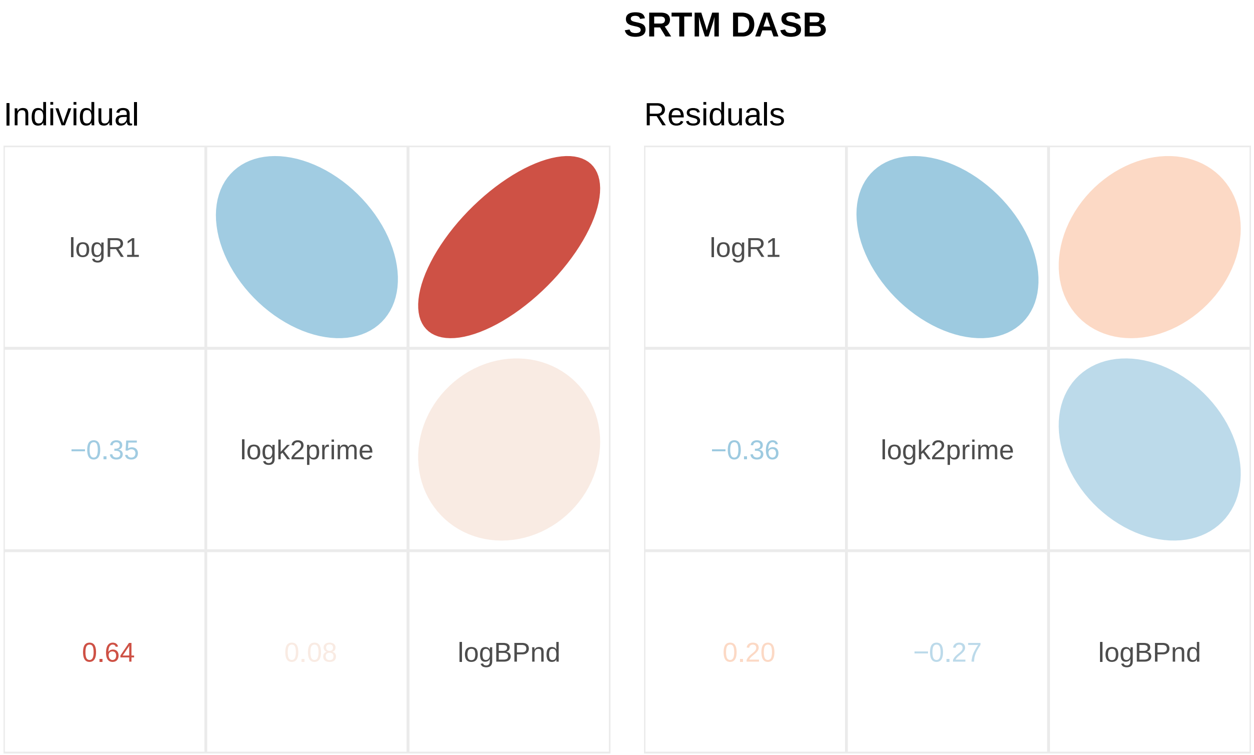

Standard Deviations

| Grouping | logR1 | logk2prime | logBPnd |
|---|---|---|---|
| Individual | 0.088 | 0.190 | 0.202 |
| Region | NA | 0.172 | NA |
| Residual | 0.068 | 0.133 | 0.176 |

$[^{11}C]$WAY100635

#### SRTM WAY

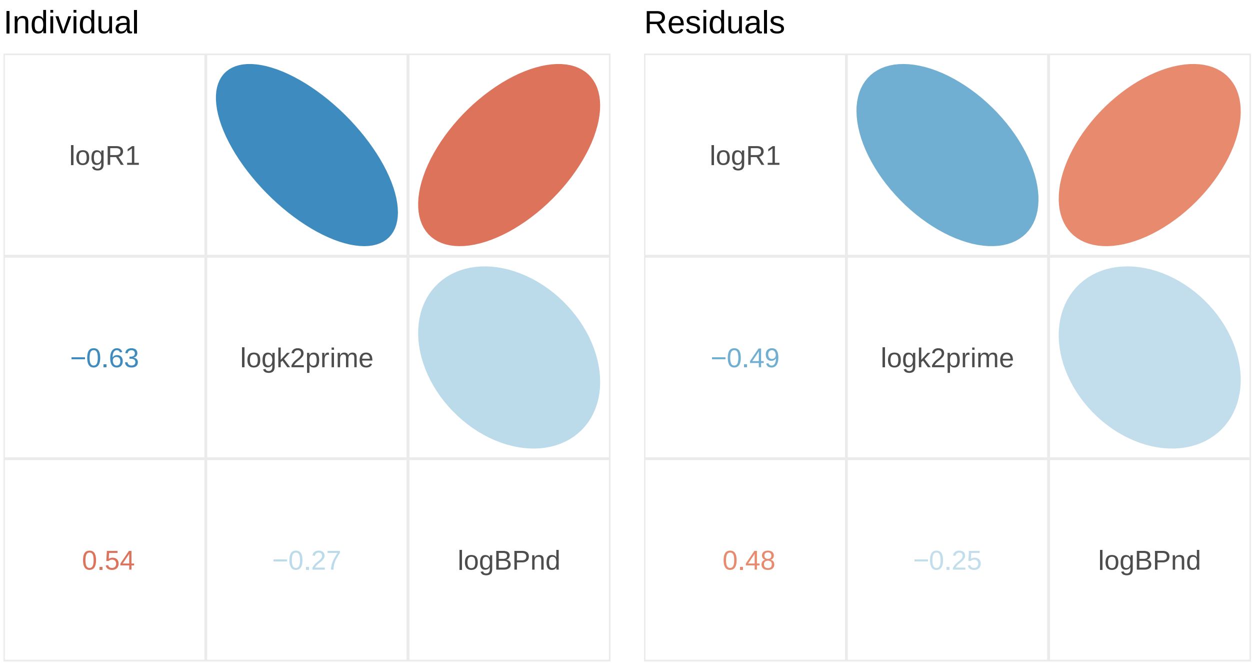

Standard Deviations

| Grouping | logR1 | logk2prime | logBPnd |
|---|---|---|---|
| Individual | 0.123 | 0.142 | 0.230 |
| Region | NA | 0.149 | NA |
| Residual | 0.059 | 0.103 | 0.127 |

### Supplementary Materials S3 : TAC Simulation Additional Figures

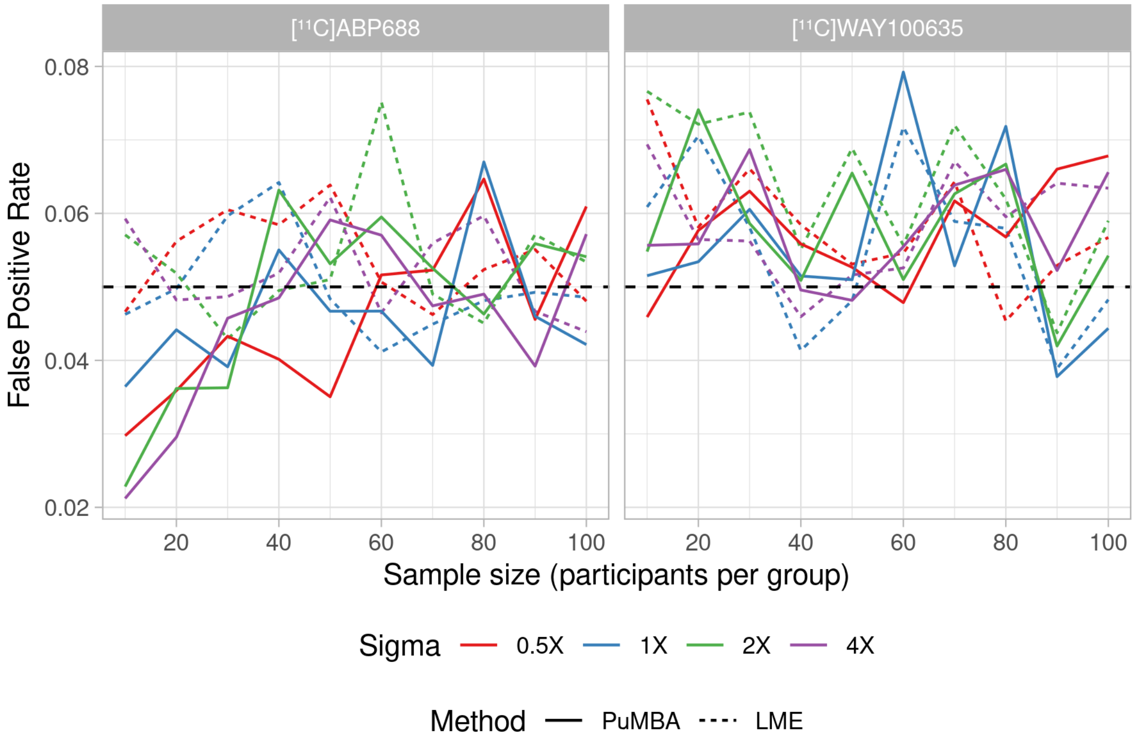

Figure 1: False positive rate across simulated datasets.

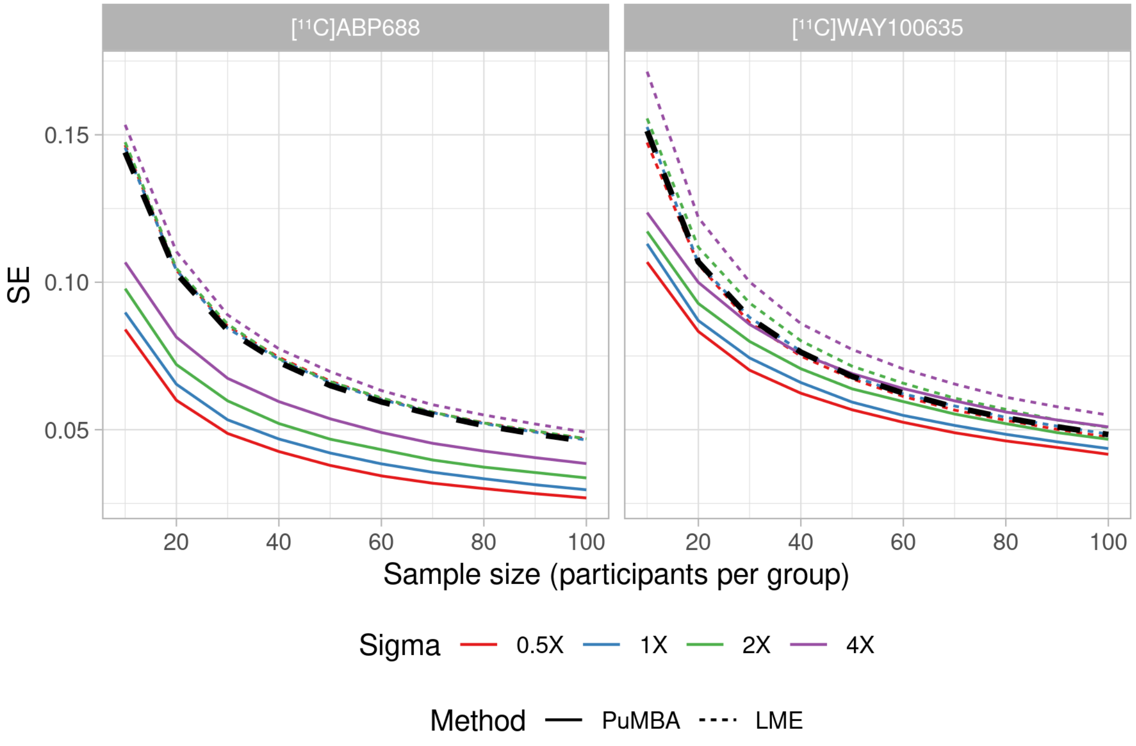

Figure 2: Mean standard error across simulated datasets.

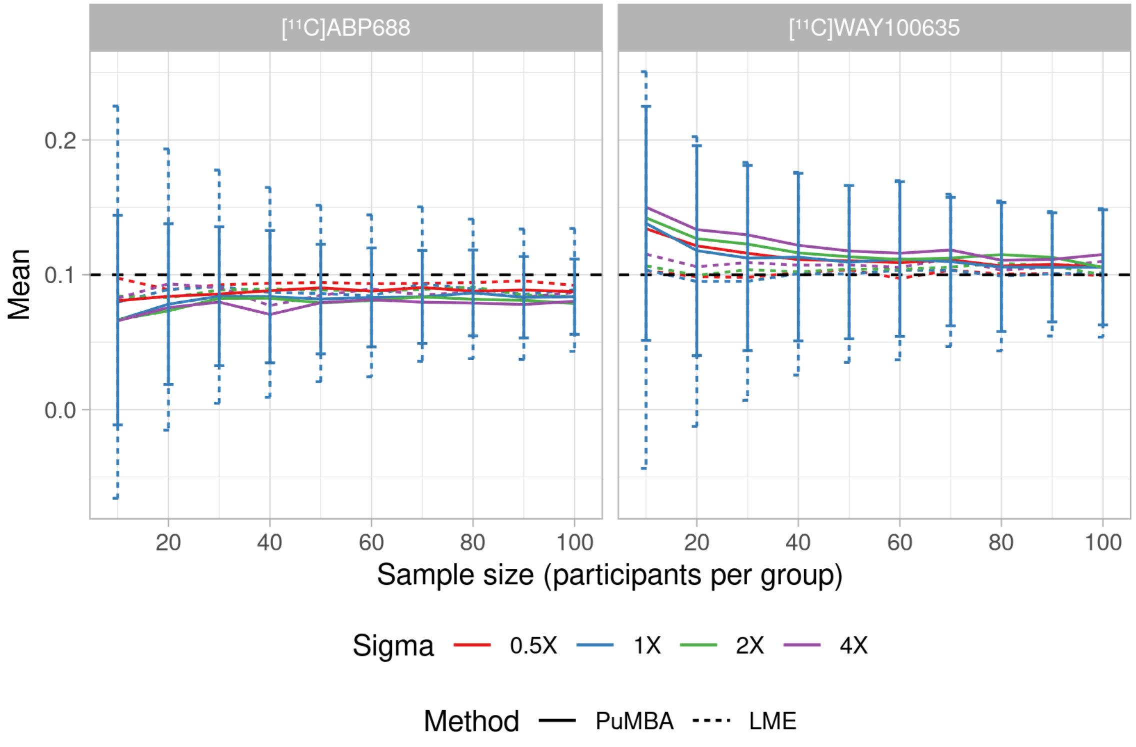

Figure 3: Mean estimated differences across simulated datasets.

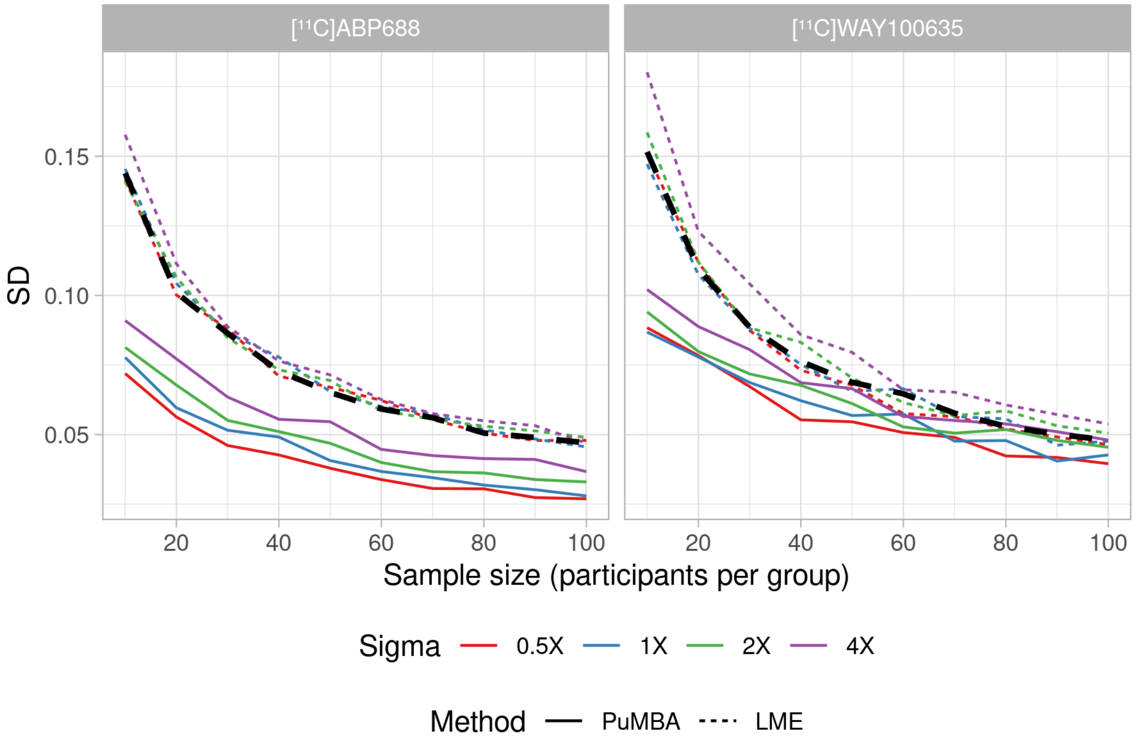

Figure 4: Standard deviation of estimated differences across simulated datasets.

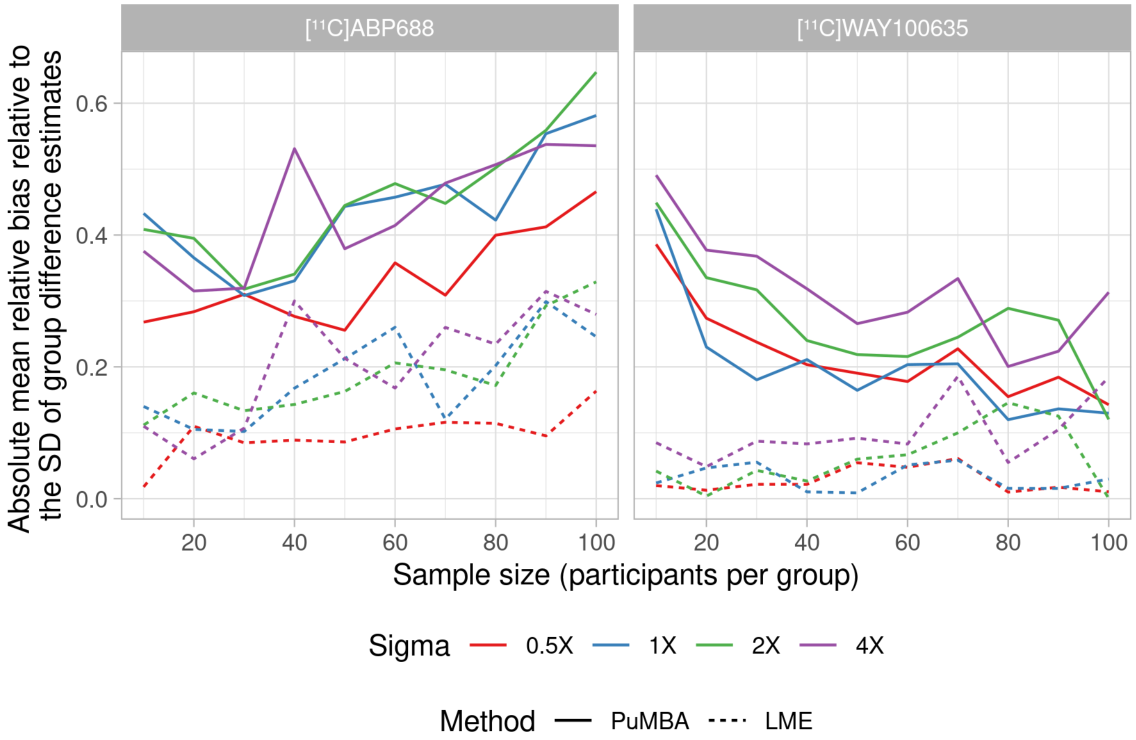

Figure 5: Bias of the estimated group difference estimates relative to the SD of estimated group differences across simulated datasets.

#### Supplementary Materials S4 : Comparison with SiMBA Additional Figures

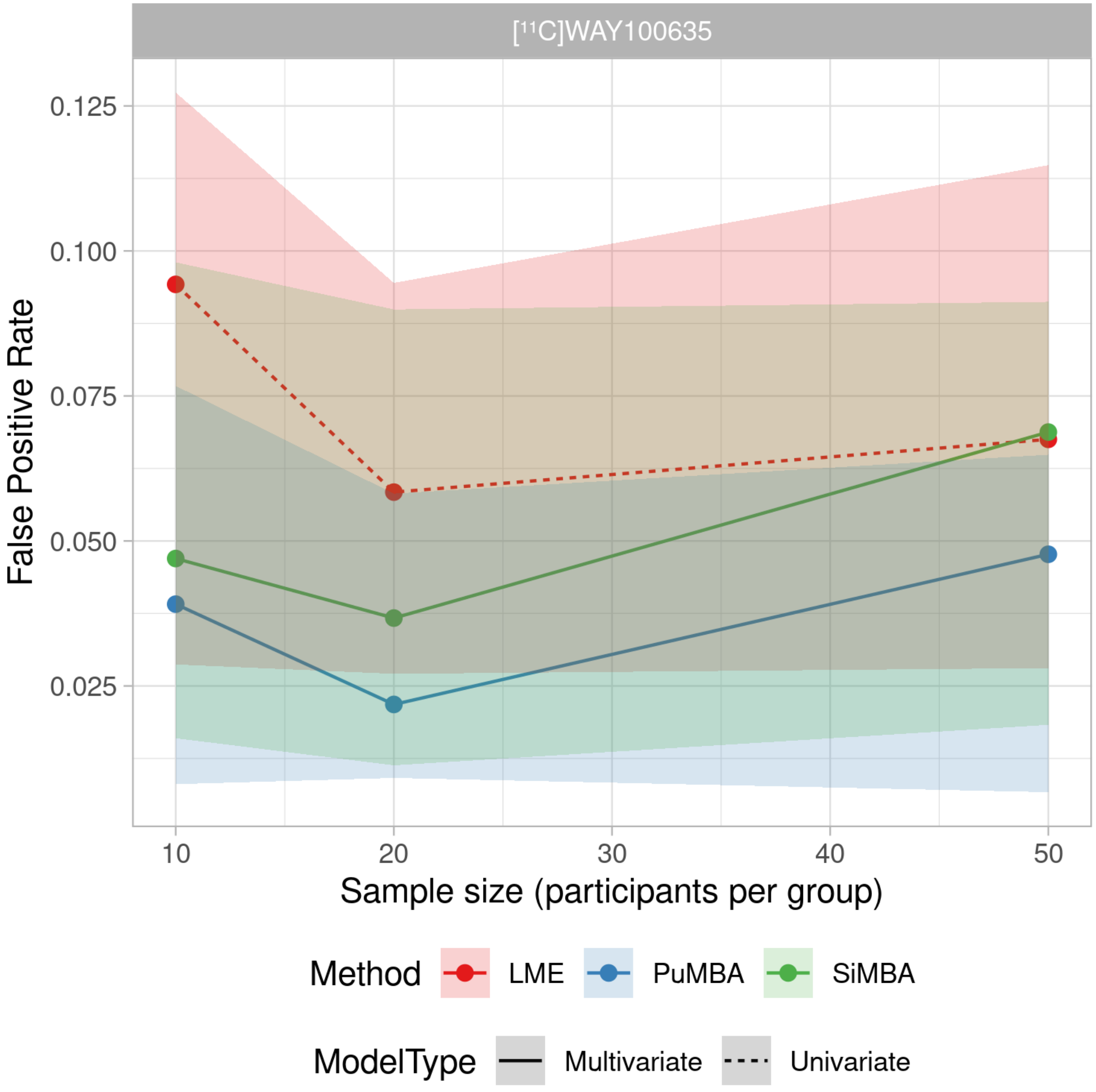

Figure 6: False positive rate across simulated datasets.

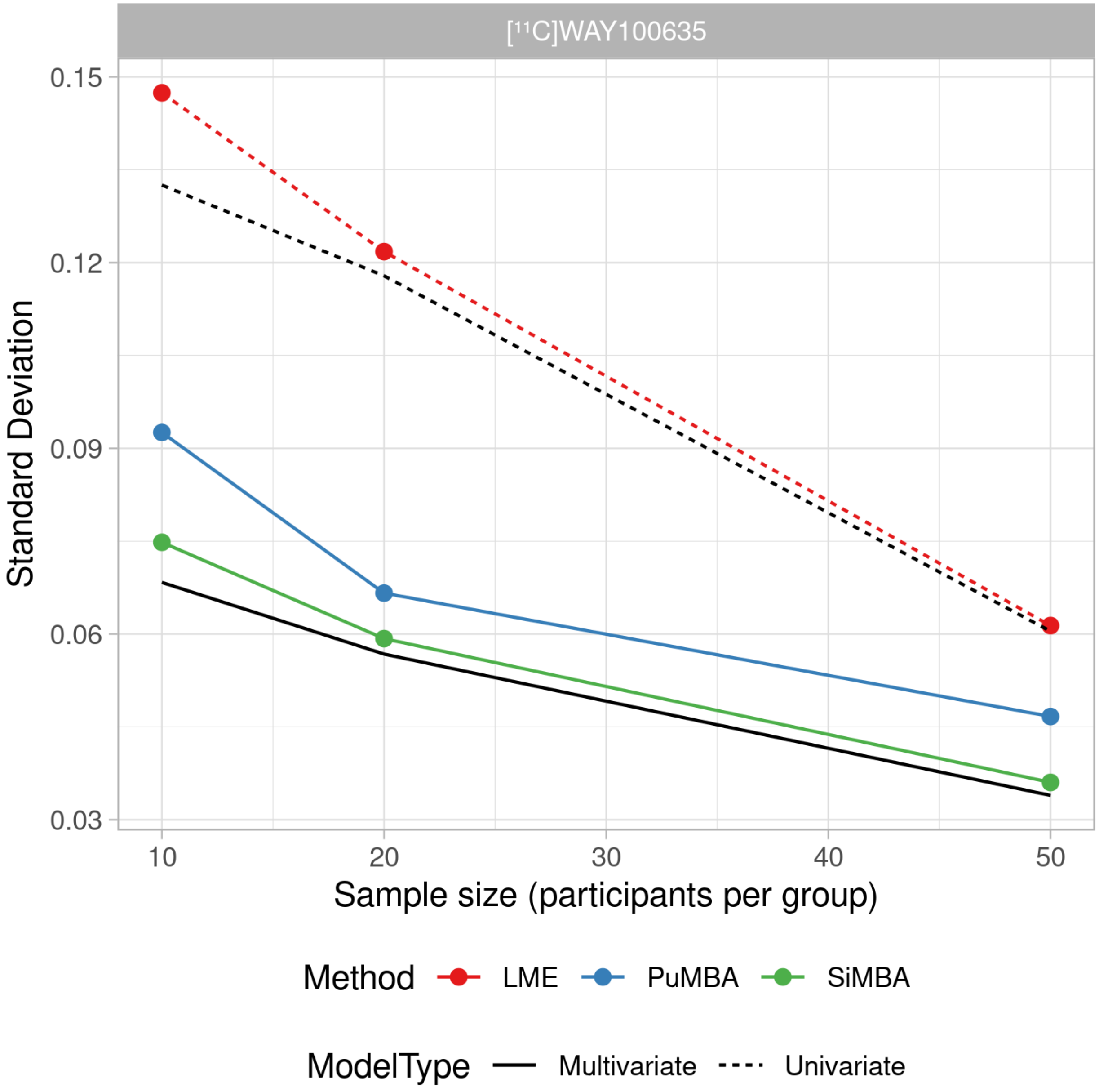

Figure 7: Standard deviation of estimated differences across simulated datasets.

### Supplementary Materials S5 : Correlated and Uncorrelated Data

To assess correlation matrix recovery, we simulated both both parameters as well as TACs with the same mean and variance for all parameters, but using univariate instead of multivariate distributions, i.e. with no correlation structure between the parameters, for the individual ($\tau$) and residual ($\epsilon$) distributions in the parameter simulations, and for the individual and individual $\times$ region distributions for the TAC simulations using SiMBA. We generated 500 simulated datasets for each condition.

For the TAC simulations, residual correlation matrices for each tracer were very similar for correlated and uncorrelated data. This suggests that the correlations in the residual variation ($\epsilon$) are primarily attributable to errors introduced during TAC fitting using NLS to generate the PK parameters owing to imperfect identifiability. Furthermore, comparing sample sizes of n=20, n=50 and n=100 per group, we observe broadly similar results, but with increased sample-to-sample variation in the smaller samples, suggesting that the poor parameter recovery is more attributable to NLS TAC fitting than to insufficient sample sizes. These residual correlation matrices differed greatly between tracers, suggesting that the nature of the imperfect identifiability is tracer-dependent.

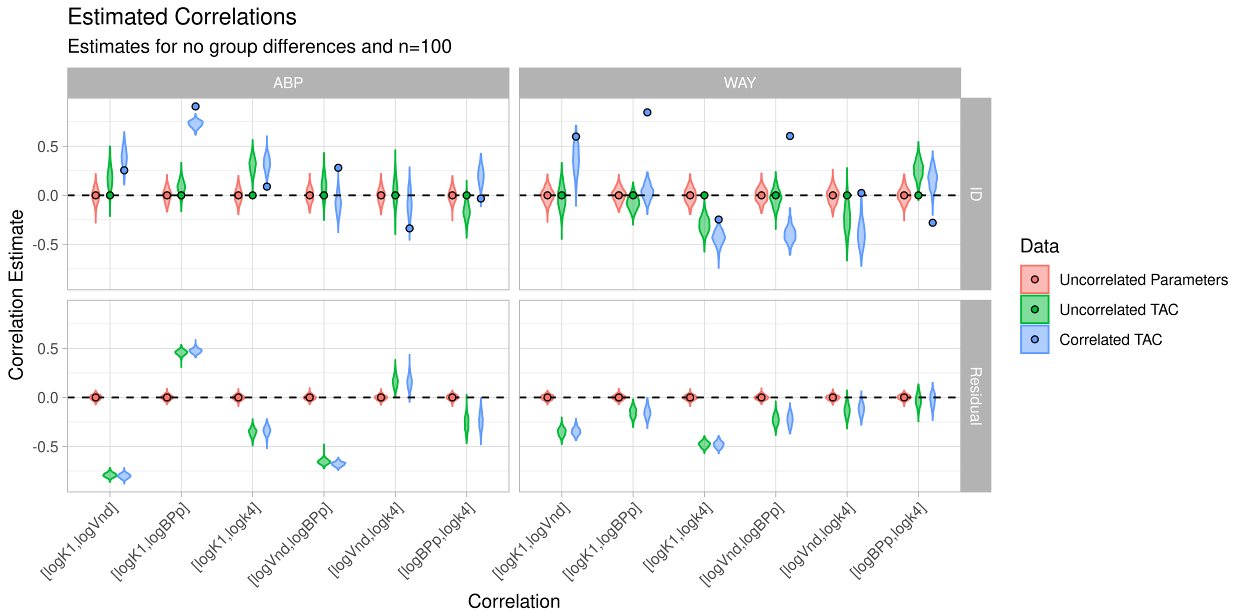

Figure 8: Estimated correlations in simulated data for sample sizes of 100 individuals per group. True values are depicted by points. Note that there no true values for the residual correlations in the TAC simulations, as the residual variance consists of a combination of individual x region variation and TAC fitting error

In the individual ($\tau$) correlation matrices, there were clear differences between the estimated correlation matrices for the correlated and uncorrelated datasets for each tracer, suggesting that the correlation structure of the data does affect the estimated multivariate correlation structure even if the recovery of the individual correlation coefficients themselves is poor. As shown below, in uncorrelated data relative to correlated data, false positive rates are unchanged, however power is reduced and standard error and standard deviation are increased.

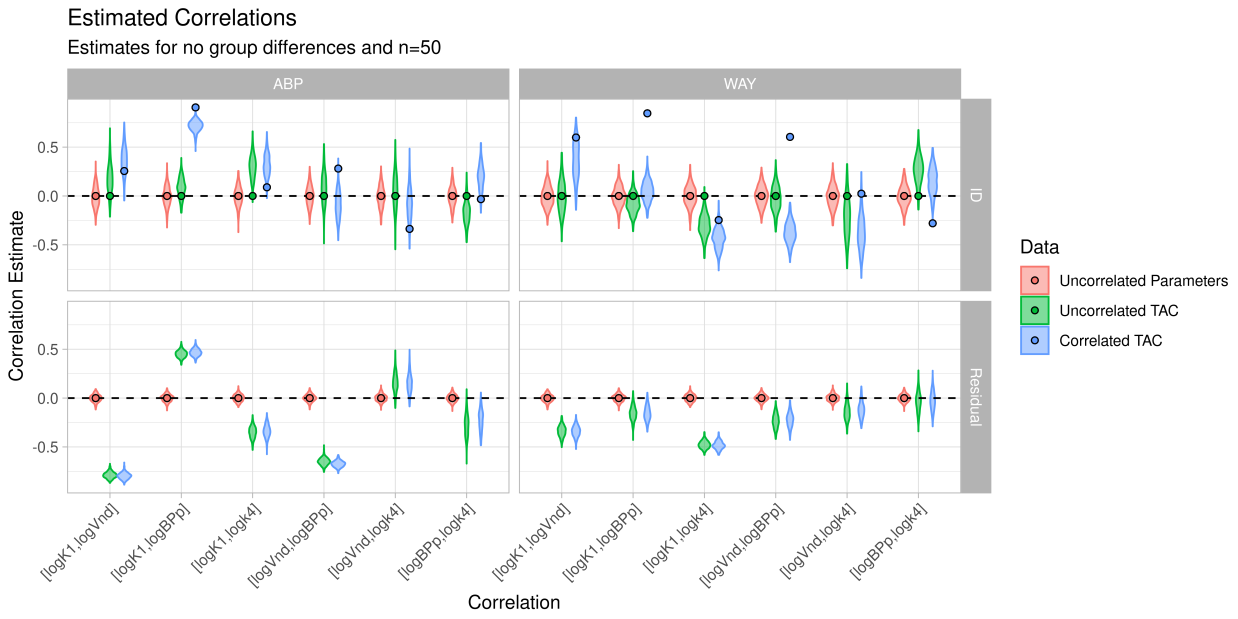

Figure 9: Estimated correlations in simulated data for sample sizes of 50 individuals per group. True values are depicted by points. Note that there no true values for the residual correlations in the TAC simulations, as the residual variance consists of a combination of individual x region variation and TAC fitting error

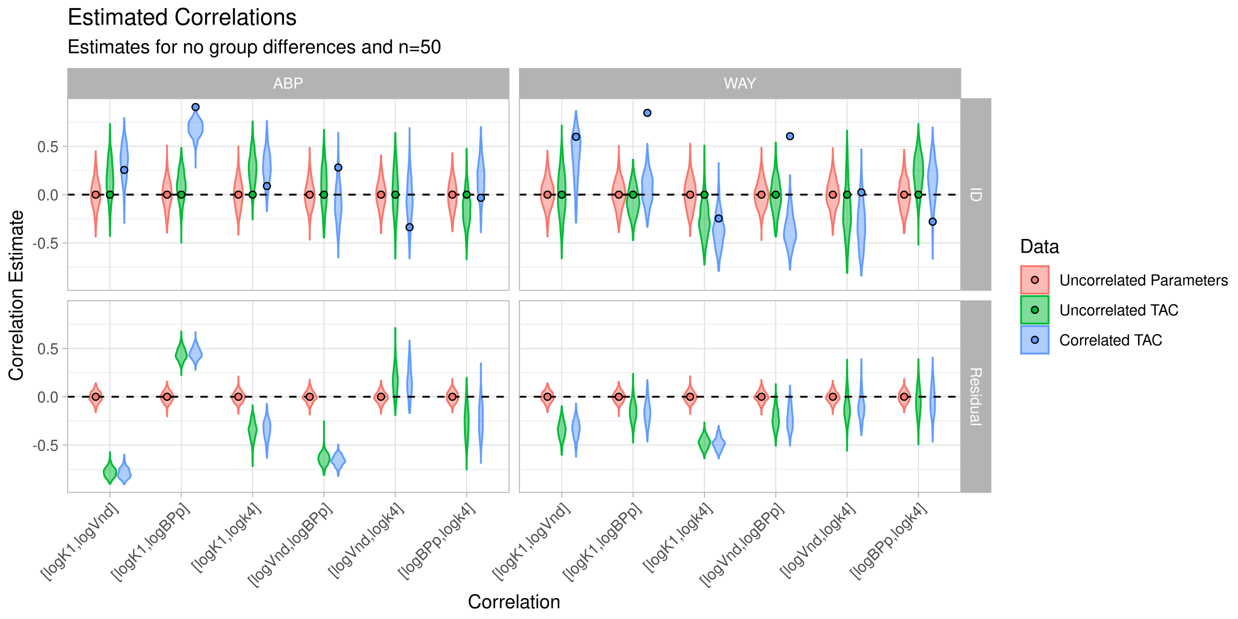

Figure 10: Estimated correlations in simulated data for sample sizes of 20 individuals per group. True values are depicted by points. Note that there no true values for the residual correlations in the TAC simulations, as the residual variance consists of a combination of individual x region variation and TAC fitting error

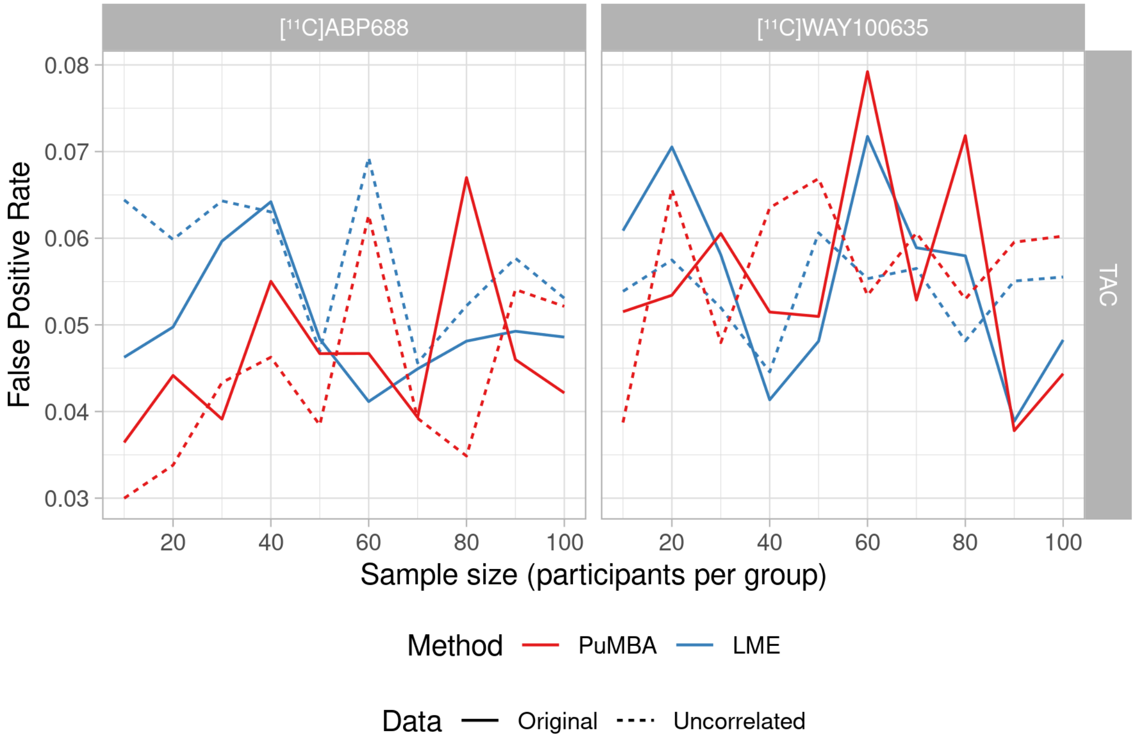

Figure 11: False positive rate across simulated datasets.

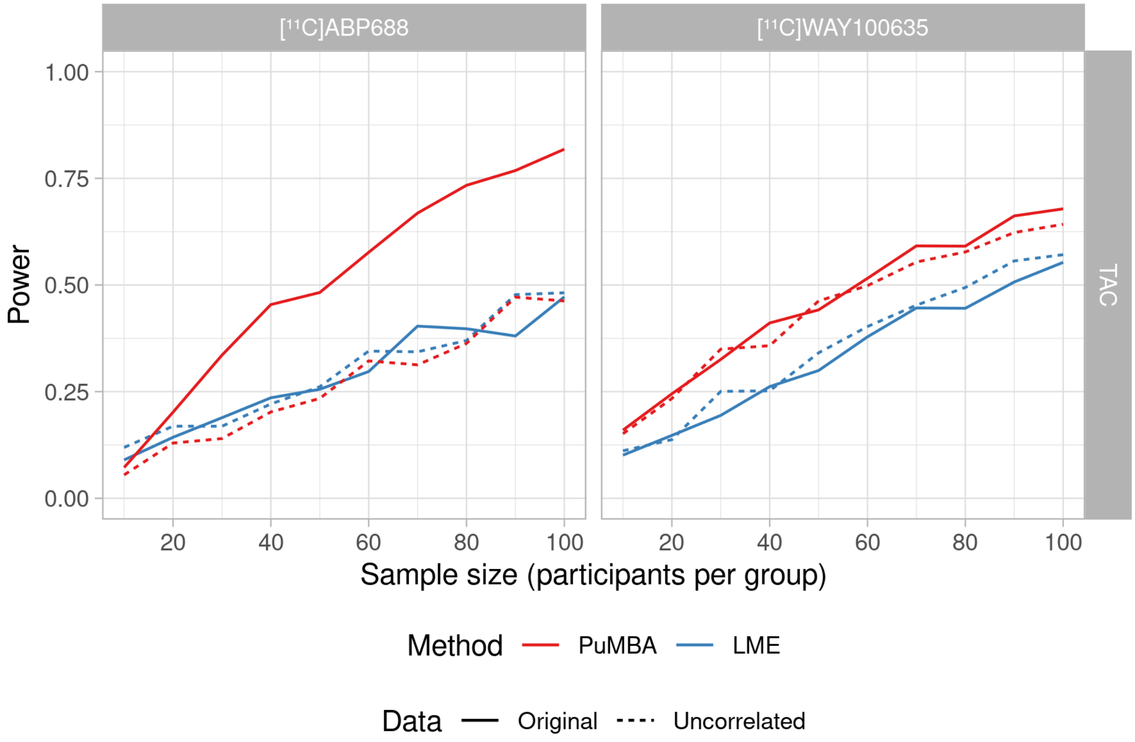

Figure 12: Power across simulated datasets.

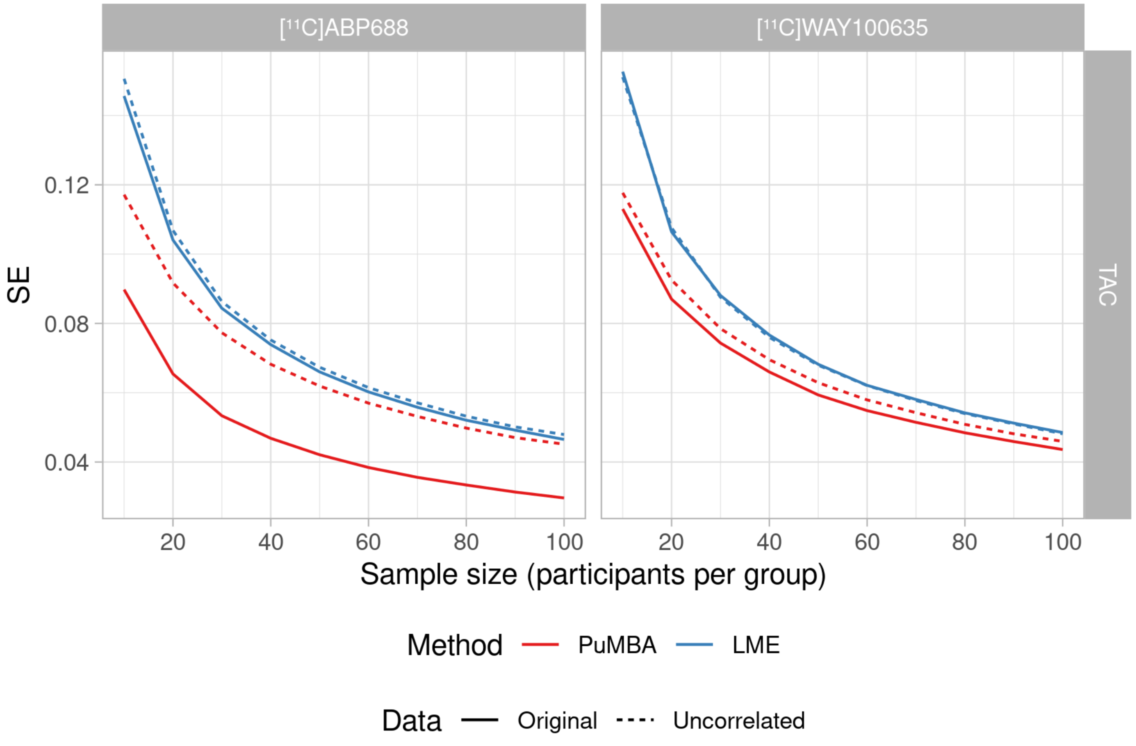

Figure 13: Mean standard error across simulated datasets.

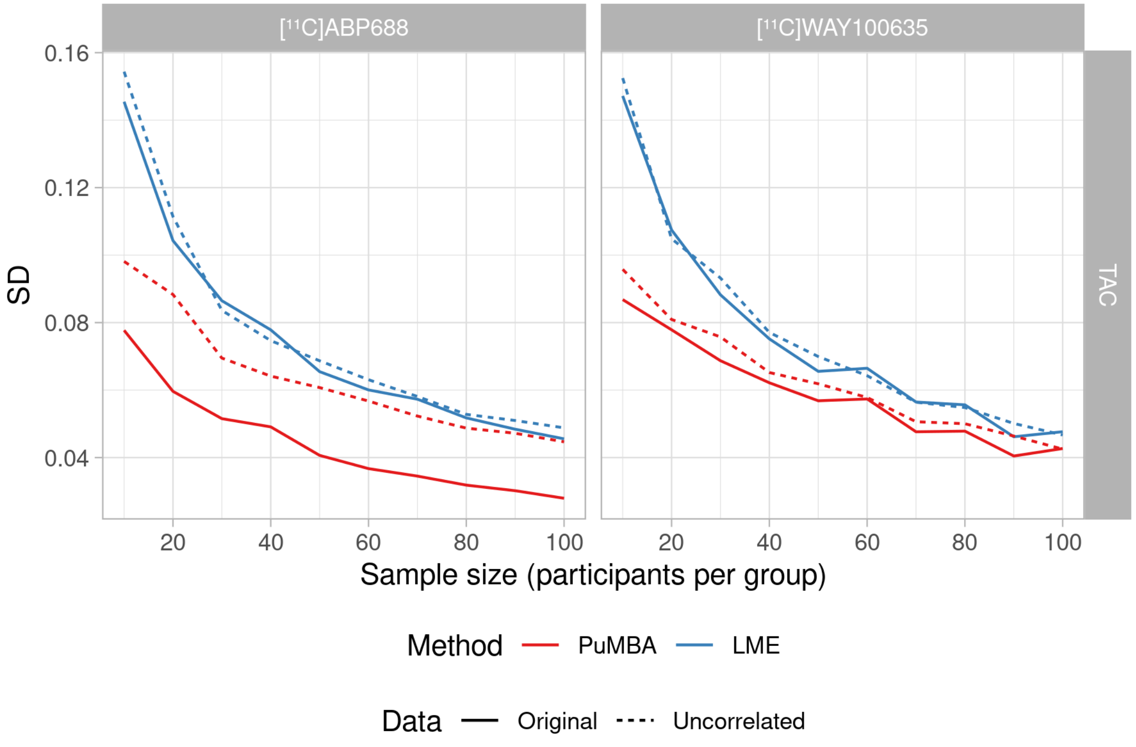

Figure 14: Standard deviation of estimated differences across simulated datasets.

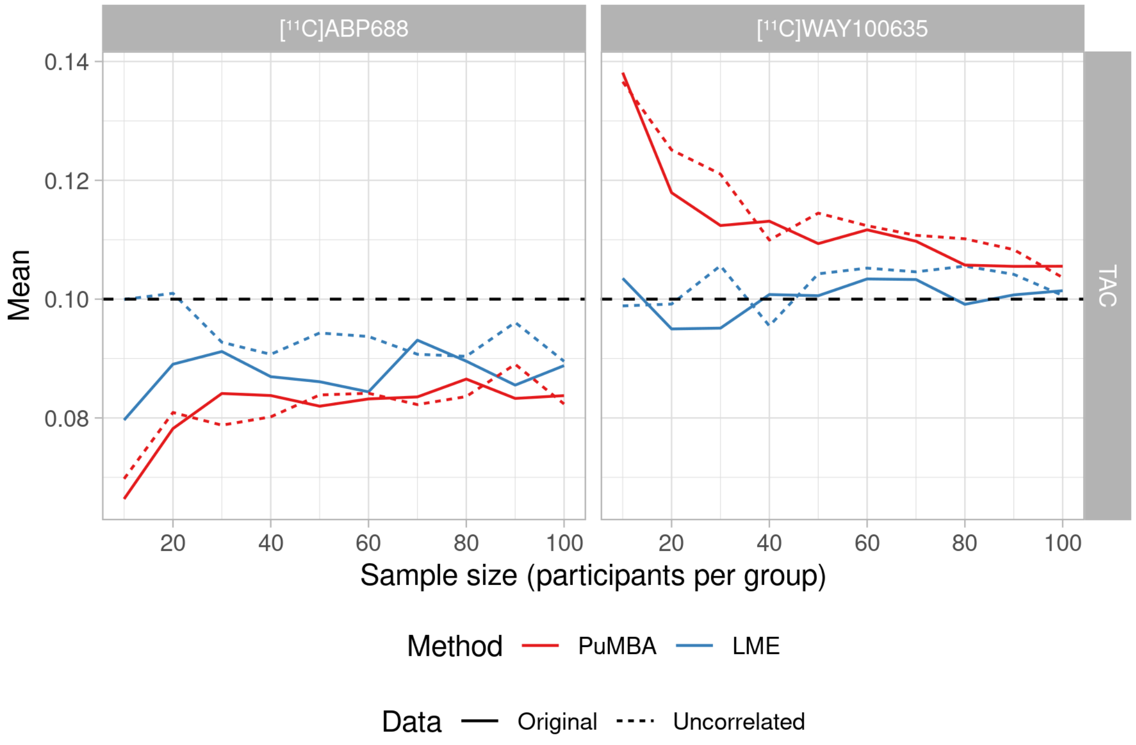

Figure 15: Mean estimated differences across simulated datasets.

#### Supplementary Materials S6 : Parameter Simulation Additional Figures

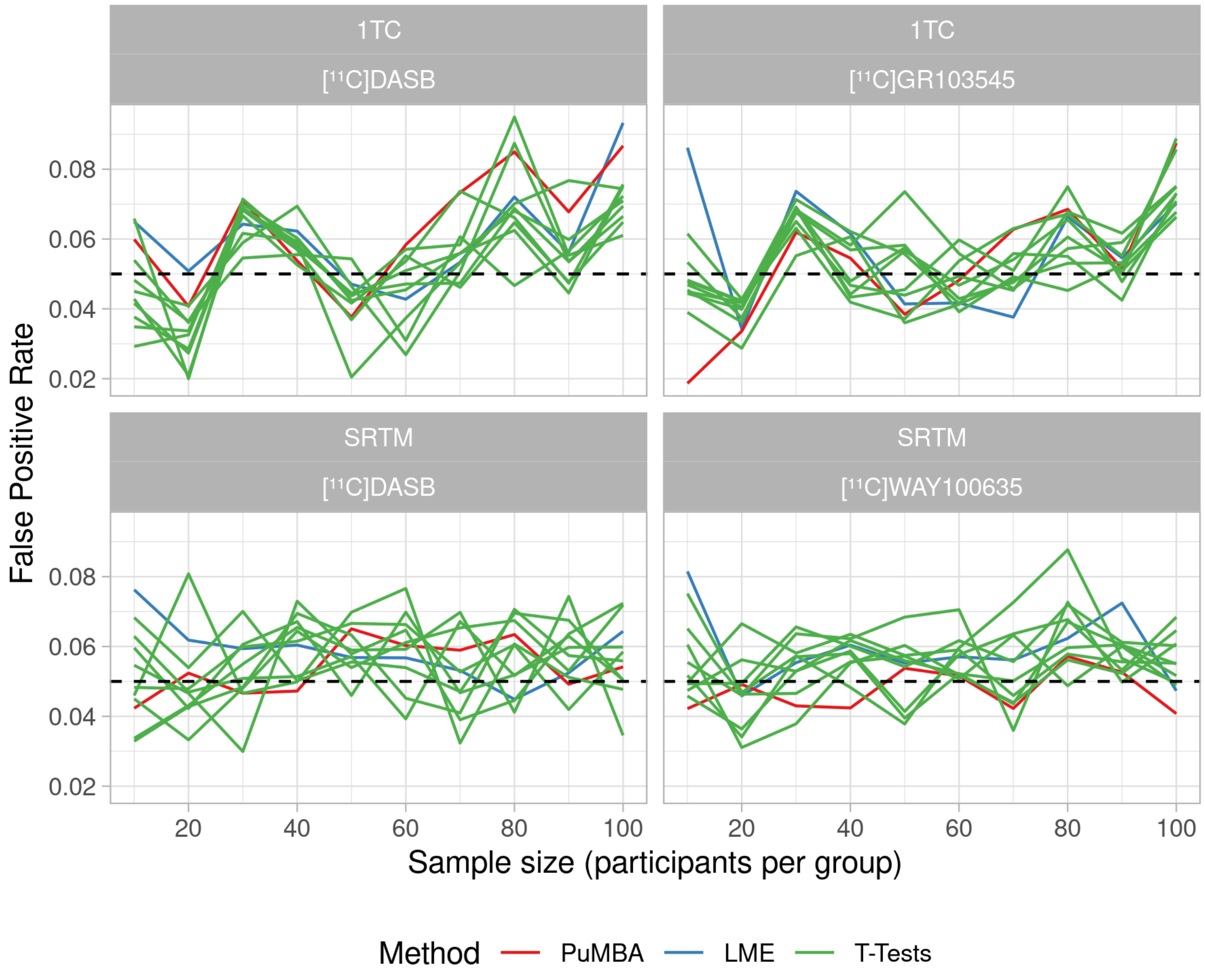

Figure 16: False positive rate across simulated datasets.

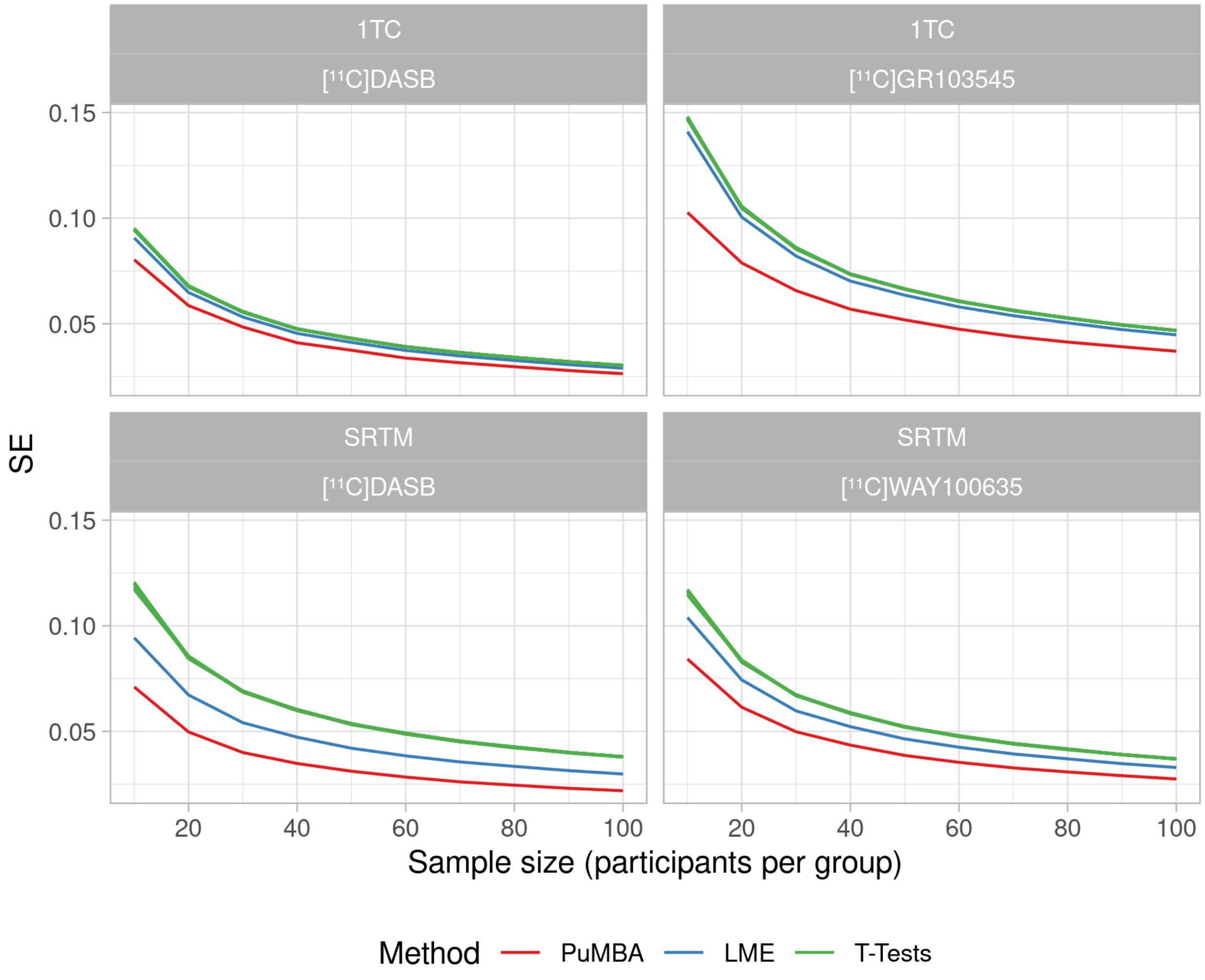

Figure 17: Mean standard error across simulated datasets.

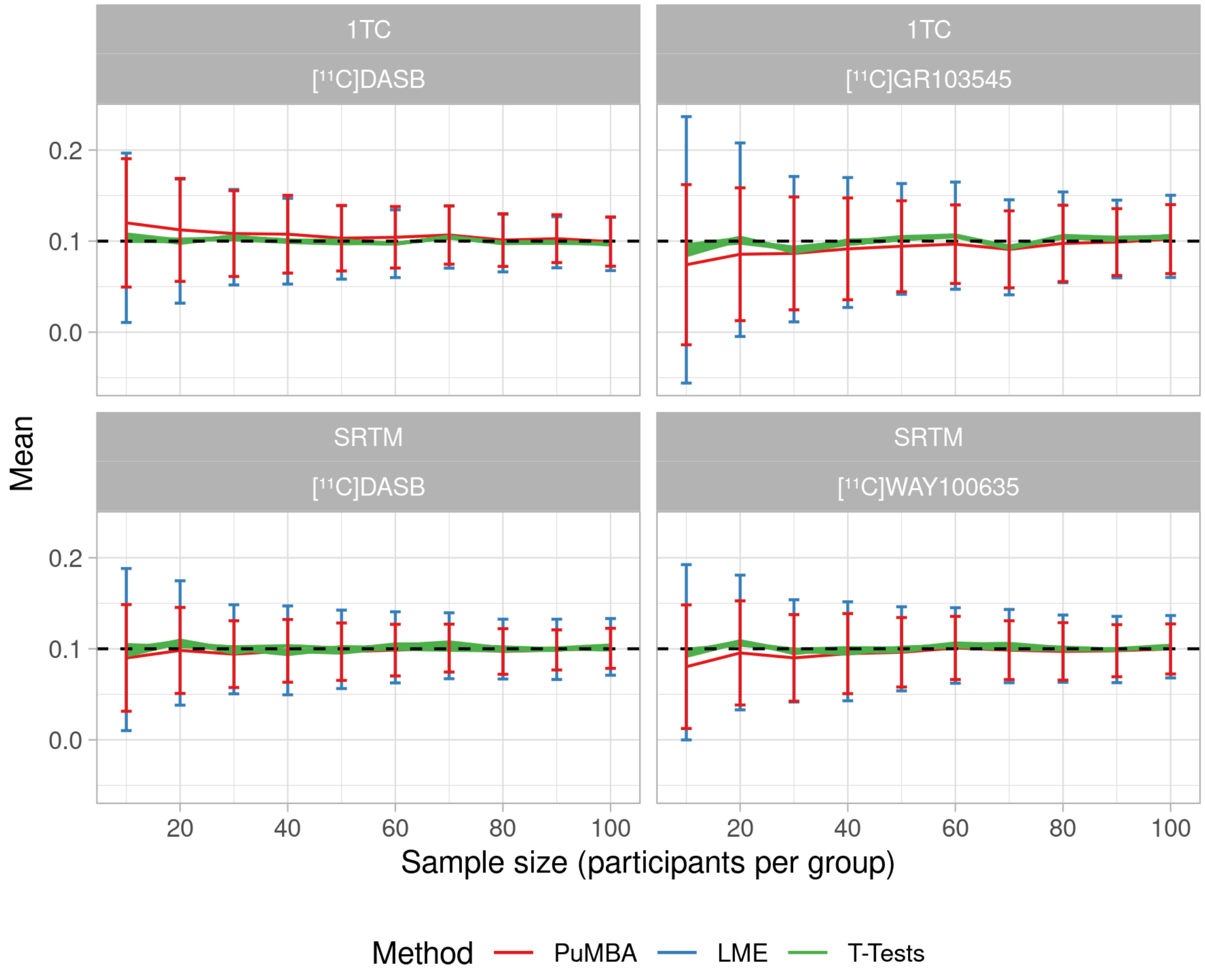

Figure 18: Mean estimated differences across simulated datasets.

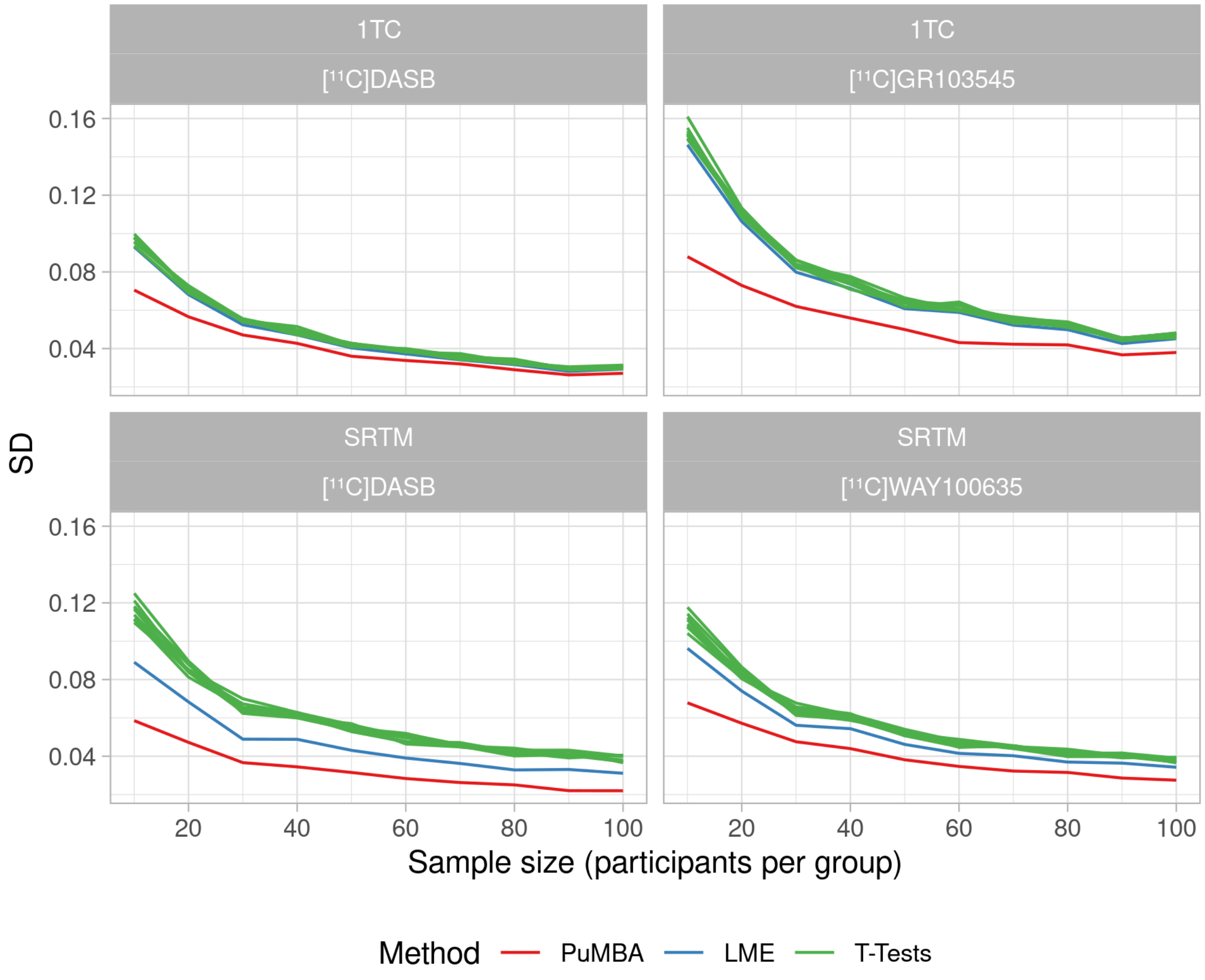

Figure 19: Standard deviation of estimated differences across simulated datasets.
